# A common SEC23B missense mutation in congenital dyserythropoietic anemia type II leads to growth restriction and symptoms of chronic pancreatitis in mice

**DOI:** 10.1101/2021.05.10.443273

**Authors:** Wei Wei, Zhigang Liu, Chao Zhang, Rami Khoriaty, Min Zhu, Bin Zhang

## Abstract

Human loss-of-function mutations in *SEC23B* result in congenital dyserythropoietic anemia type II (CDAII). Complete deficiency of SEC23B in mice leads to perinatal death caused by massive degeneration of professional secretory tissues. Functions of SEC23B in postnatal mice are unclear. In this study, we generated knockout (KO) mice with deletion of exons 5 and 6 of *Sec23b* and knockin (KI) mice with the E109K mutation, the most common CDAII missense mutation. Despite decreased SEC23B protein level, *Sec23b^ki/ki^* mice showed no obvious abnormalities. Hemizygous (*Sec23b^ki/ko^*) mice exhibit a partial lethal phenotype, with only half of expected mice survive past weaning. *Sec23b^ki/ko^* pancreas had exocrine insufficiency and chronic pancreatitis histology, as well as increased ER stress and apoptosis. Moreover, *Sec23b^ki/ko^* mice exhibited severe growth restriction accompanied by growth hormone (GH) insensitivity, reminiscent of the Laron syndrome. Growth restriction is not associated with hepatocyte-specific *Sec23b* deletion, suggesting non-liver origin of the phenotype. Inflammation associated with chronic pancreatic deficiency may explain GH insensitivity in *Sec23b^ki/ko^* mice. Our results indicate that phenotype severities are linked to the residual functions of SEC23B, demonstrating the importance of SEC23B in pancreatic acinar function in adult mice. The *Sec23b^ki/ko^* mice provide a novel model of chronic pancreatitis and growth retardation.

## Introduction

Coat protein complex II (COPII) vesicles transport approximately 8,000 mammalian proteins from the endoplasmic reticulum (ER) to the Golgi apparatus (1–4). COPII is composed of five core components, including SAR1, SEC23, SEC24, SEC13 and SEC31 (5, 6). COPII vesicles assembly begins at distinct ER exit sites on the cytosolic surface marked by SEC16, where GDP-bound SAR1 is converted to GTP-bound SAR1 by the GTP exchange protein SEC12. SAR1-GTP recruits the SEC23-SEC24 heterodimer to form the inner layer of the COPII coat (7–9). The SAR1-SEC23-SEC24 prebudding complex further recruits the outer layer composed of SEC13-SEC31 heterotetramers, facilitating the budding of COPII vesicles from the surface of the ER (10–12).

In contrast to yeast, mammalian genomes contain two or more paralogues for most genes encoding COPII components (4, 13). Among these are two SEC23 paralogues, SEC23A and SEC23B, which share ~85% amino acid sequence identity (4, 13). In humans, missense mutations in *SEC23A* lead to cranio-lenticulo-sutural dysplasia, characterized by craniofacial and skeletal abnormalities, in part due to collagen secretion defects (14, 15). We previously showed that mice with SEC23A deficiency exhibit mid-embryonic lethality, defective development of extraembryonic membranes and neural tube opening in the midbrain, associated with secretion defects of multiple collagen types (16). Human mutations in *SEC23B* result in congenital dyserythropoietic anemia type II (CDAII) (17, 18), characterized by mild to moderate anemia, bi/multinucleated erythroblasts in the bone marrow, double membrane appearance in the red blood cell, and a faster shift and narrower band of RBC membrane protein band 3 on SDS-PAGE (19, 20). In CDAII patients, the majority of mutations are missense mutations, and no patients with two null mutations have been reported (21). E109K and R14W are two of the most frequent recurrent mutations in CDAII patients (22). Patients that are compound heterozygous with a null mutation and a missense mutation tend to have more severe anemia phenotypes than patients with two missense mutations (23), and hypomorphic mutations account for mild phenotypes of CDAII (24). In addition to causing CDAII, SEC23B mutations are also linked to Cowden’s syndrome (25).

Despite the identification of genetic defects, the molecular mechanism of CDAII caused by SEC23B deficiency in humans remains unknown. We previously reported that mice with near complete deficiency for SEC23B were born with no apparent anemia phenotype, but died shortly after birth, with degeneration of professional secretory tissues, in particular degeneration of the pancreas (26). Pancreatic deficiency of SEC23B is the cause of perinatal lethality and SEC23B is essential for the normal function of pancreatic acinar cells in adult mice (27, 28). Hematopoietic deficiency of SEC23B does not result in CDAII phenotype in adult mice (29). A *Sec23a* coding sequence inserted into the murine *Sec23b* locus completely rescues the lethal SEC23B-deficient pancreatic phenotype (30). However, it is unknown whether murine SEC23B is critical for functions outside of professional secretory tissues. This question has been hampered by the lack of a global SEC23B deficient mouse model that can survive beyond the first day after birth.

Here we report the phenotype of SEC23B deficient mice with the E109K missense mutation, which survive to adulthood. E109K hemizygous mice exhibited severe postnatal growth retardation accompanied by chronic pancreatitis. Our results suggest that global SEC23B deficiency is not associated with a CDAII-like phenotype in mice. SEC23B plays a critical role in normal mouse postnatal growth and pancreatic functions.

## Results

### The E109K mutation results in decreased SEC23B protein level

E109K is a founder mutation in the Italian population and also the most common mutation reported in CDAII patients (22). To study the impact of the E109K mutation on SEC23B function *in vivo*, we generated two mouse lines. One is a SEC23B knockout (KO) mouse line with deletion of exons 5 and 6 (Fig. 1A and 1B, Fig. S1). This mouse is distinct from the previously reported conditional gene-trap mouse that also deleted exons 5 and 6 (29). The second is a knockin (KI) mouse line carrying the SEC23B^E109K^ mutation (Fig. 1C and 1D, Fig. S2). Intercrosses of heterozygous KO (*Sec23b^ko/+^*) mice produced homozygous KO (*Sec23b^ko/ko^*) pups that die shortly after birth with pancreas degeneration (Fig. S3), consistent with previously reported phenotype of SEC23B-deficient mice. Intercrosses of heterozygous KI (*Sec23b^ki/+^*) mice produced homozygous KI (*Sec23b^ki/ki^*) mice. Intercrosses of heterozygous knockout (*Sec23b^ko/+^*) mice with *Sec23b^ki/ki^* mice generated mice hemizygous for the SEC23B^E109K^ mutation (*Sec23b^ki/ko^*). Levels of SEC23A, SEC23B and total SEC23 proteins from WT, *Sec23b^ki/ki^* and *Sec23b^ki/ko^* mice were measured in liver, pancreas and salivary gland lysates by immunoblotting. As shown in Fig. 1E, SEC23B levels in *Sec23b^ki/ki^* mice decreased in all the tested organs compared to WT mice. SEC23B protein levels further decreased in *Sec23b^ki/ko^* mice compared to those of *Sec23b^ki/ki^* mice. Compensatory increase in SEC23A levels had been reported in tissues of SEC23B knockout mice (26). We observed that SEC23A levels increased in both *Sec23b^ki/ki^* and *Sec23b^ki/ko^* mice compared to that of WT mice (Fig. 1E). Increased SEC23A levels were sufficient to offset the decreased SEC23B, so that steady-state total SEC23 levels were comparable between WT and *Sec23b^ki/ki^* mouse tissues tested. However, increased SEC23A levels were insufficient to offset decreased SEC23B levels in *Sec23b^ki/ko^* mice, so that lower steady-state levels of total SEC23 were detected in *Sec23b^ki/ko^* mouse tissues (Fig. 1E). In WT mouse pancreas, the SEC23B level gradually decreased as mice age, whereas the SEC23A level gradually increased. However, in *Sec23b^ki/ko^* mouse pancreas, levels of both SEC23 paralogs remain relatively steady (Fig. S4).

**Figure 1.**
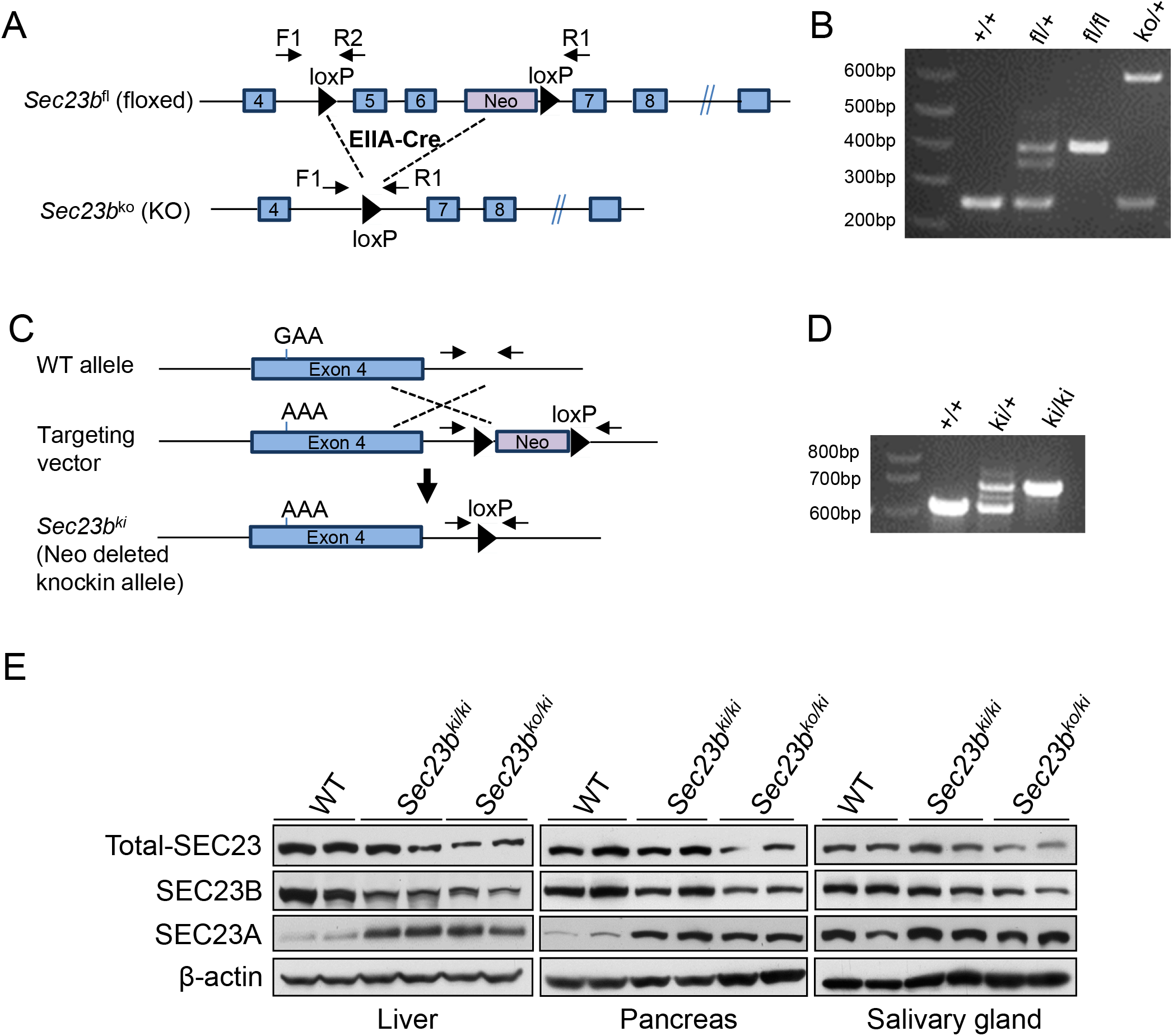
Generation of *Sec23b* knockout and knockin alleles. A) Schematics of the *Sec23b* floxed allele and the conditional knockout allele with deletion of exons 5 and 6. Positions of primer are indicated. B) A three-primer PCR assay distinguished floxed allele, knockout allele, and WT allele. C) Schematics of the knockin allele with the E109K mutation within the exon 4. D) A two-primer PCR assay distinguished knockin and WT alleles. Detailed diagrams on the generation of KO and KI alleles are shown in Fig. S1 and Fig. S2. E) Immunoblotting analysis of SEC23B, SEC23A and total SEC23 in liver, pancreas and salivary glands from 2-month old mice of the indicated genotypes. For each genotype, tissues from 2 mice were analyzed.

We also stably expressed human WT SEC23B, SEC23B^E109K^ and SEC23B^R14W^ mutations as GFP fusion proteins in Nthy-ori 3-1 cells. The R14W mutation is another founder mutation in the Italian population and the second most common mutation reported in CDAII patients (22). Consistent with the mouse results, we observed much lower expression of SEC23B^E109K^ and SEC23B^R14W^ than the WT protein (Fig. S5A). Expression of WT SEC23B, but not SEC23B mutants, led to a decrease in the SEC23A level (Fig. S5A). Both GFP-tagged WT SEC23B and SEC23B^R14W^ proteins co-localized with SEC16A, a marker for the ER exit sites (Fig. S5B). However, GFP-tagged SEC23B^E109K^ mutant failed to co-localize with SEC16A and became evenly distributed in the cytosol (Fig. S5B), suggesting that E109K and R14W mutants have different defects. Due to the lack of an antibody suitable for immunofluorescence, we were unable to determine the subcellular localization pattern of SEC23B^E109K^ in KO mouse cells.

### Sec23b^E129K^ hemizygosity leads to partial lethality and growth restrictions in the mouse

At the time of weaning, the number of *Sec23b^ki/ki^* mice was slightly lower than expected, although the genotype distribution is not statistically significantly different from the expected (P>0.16, Table 1). At normal weaning time (day 21), the observed number of *Sec23b^ki/ko^* mice were significantly lower than expected (P<0.001, Table 2). To determine whether the loss of *Sec23b^ki/ko^* pups occurred postnatally, we genotyped neonatal pups from the *Sec23b^ko/+^* X *Sec23b^ki/ki^* cross and found no significant loss of *Sec23b^ki/ko^* neonates (P>0.41, Table 2), suggesting that the loss of *Sec23b^ki/ko^* pups primarily occurred postnatally. Heterozygous *Sec23b^ki/+^* and *Sec23b^ko/+^* mice exhibited normal survival, and no abnormalities were noted during routine necropsy.

**Table 1.** Genotype distribution of pups at weaning from intercrosses of *Sec23b^ki/+^* mice. Chi-square test was used to calculate the P value for the real ratio of *Sec23b*^ki/ki^ versus expected ratio of *Sec23b*^ki/ki^ mice

|  | Genotype |  |  | P value |
| --- | --- | --- | --- | --- |
|  | +/+ (n) | ki/+ (n) | ki/ki (n) |  |
| Expected ratio | 25 | 50 | 25 |  |
| Observed ratio (n=195) | 28.7 (56) | 51.8 (101) | 19.5 (38) | >0.16 |

**Table 2.** Genotype distribution of pups from intercrosses of *Sec23b^ki/ki^* mice with *Sec23b*^ko/+^ mice. Chi-square test was used to calculate the P value for real ratio of *Sec23b^ki/ko^* versus expected ratio of *Sec23b^ki/ko^* mice

|  | Genotype |  | P value |
| --- | --- | --- | --- |
|  | ki/+ (n) | ki/ko (n) |  |
| Expected ratio | 50% | 50% |  |
| P21 Observed ratio (n=190) | 75.6% (144) | 24.4% (46) | <0.001 |
| P1 Observed ratio (n=98) | 54% (53) | 41% (45) | >0.41 |

We noticed that *Sec23b^ki/ko^* mice appeared smaller in size compared to WT controls (Fig. 2A). In contrast, *Sec23b^ki/ki^* and *Sec23b^ki/+^* mice are indistinguishable from WT controls. Therefore, we monitored the body weight and body length of WT, *Sec23b^ki/ki^* and *Sec23b^ki/ko^* mice from the ages of 2 weeks to 18 weeks. As shown in Fig. 2B, the body weight of both male and female *Sec23b^ki/ko^* mice were consistently lower than that of WT or *Sec23b^ki/ki^* mice throughout the study period. The differences among these mouse groups are especially striking within the first 6 weeks of life, with the body weights of *Sec23b^ki/ko^* mice reaching only 1/3 of WT and *Sec23b^ki/ki^* mice at week 4 (Fig. 2B). The differences narrowed between 8 and 12 weeks, and remained relatively steady afterwards. The trend of body length is very similar to that of body weight (Fig. 2C), which showed significantly decreased body length of *Sec23b^ki/ko^* mice compared to the two other groups of mice. Even though an apparent growth spurt occurred in *Sec23b^ki/ko^* mice from 7 to 10 weeks, both male and female adult *Sec23b^ki/ko^* mice remain significantly shorter than WT and *Sec23b^ki/ki^* mice. In contrast to *Sec23b^ki/ko^* mice, the growth curve of *Sec23b^ki/ki^* mice is similar to WT controls. To determine whether the size differences existed at birth, we further measured the weight and length of the offspring of the *Sec23b^ko/+^* X *Sec23b^ki/ki^* cross as neonates (P0) and at one week after birth (P7).

**Figure 2.**
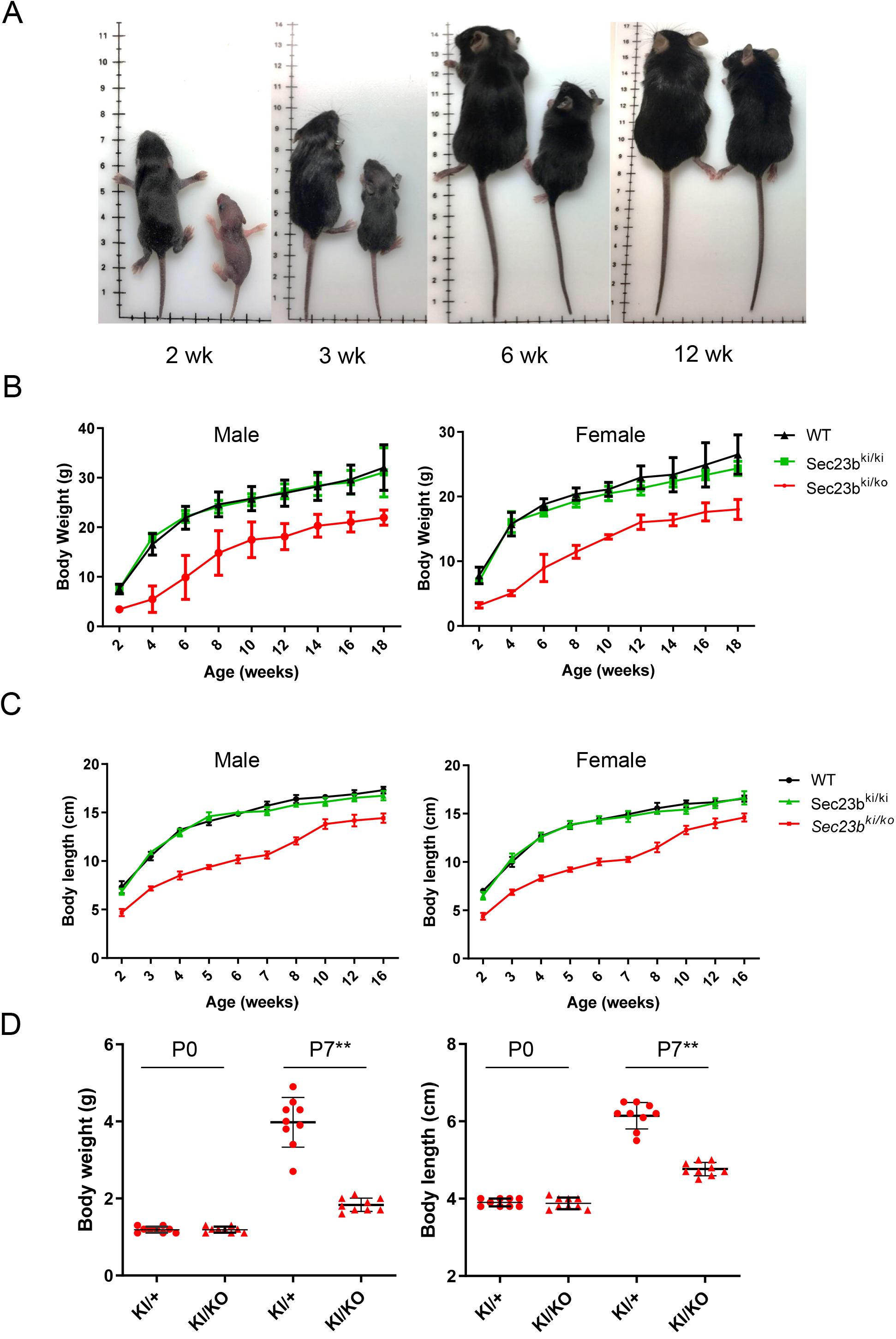
Growth restriction in *Sec23B^ki/ko^* mice. A) *Sec23b^ki/ko^* mice are consistently smaller than WT controls of the same age. B) Body weights of both male and female WT, *Sec23b^ki/ki^* and *Sec23b^ki/ko^* mice from 2 weeks of age to 18 weeks of age (data are mean ± SD, P<0.05 at all time points). C) Body lengths of both male and female WT, *Sec23b^ki/ki^* and *Sec23b^ki/ko^* mice from 2 weeks of age to 16 weeks of age (data are mean ± SD, P<0.05 at all time points). D) Body weights and lengths of *Sec23b^ki/+^* and *Sec23b^ki/ko^* pups at P0 and P7.

Although no significant differences were observed between *Sec23b^ki/+^* and *Sec23b^ki/ko^* at P0, *Sec23b^ki/ko^* pups had become significantly smaller at P7, both in weight and in length (Fig. 2D), indicating that the growth restriction occurred postnatally.

### The E109K mutation does not lead to a CDAII-like phenotype in mice

Previous studies demonstrated that neither neonates of global *Sec23b* KO mice nor adult hematopoietic cell-specific *Sec23b* KO mice showed a CDAII-like phenotype (26) (29). We performed complete blood counts of 1-month and 4-month old *Sec23b^ki/+^* and *Sec23b^ki/ko^* mice from the offspring of the *Sec23b^ko/+^* X *Sec23b^ki/ki^* cross. The red blood cell (RBC) count, hemoglobin level and hematocrit level are all significantly decreased in *Sec23b^ki/ko^* mice, although the differences have narrowed from 1 month to 4 months (Fig. 3A-B). Narrow band size and a shift in the mobility of membrane protein band 3 on SDS-PAGE are typical red blood cell abnormalities in CDAII patients. However, RBC from *Sec23b^ki/ko^* mice exhibited no abnormalities in Band 3 compared to WT RBC (Fig. 3C). A characteristic feature of bone marrow morphology of CDAII patients is the increased number of binuclear erythroblasts. However, no binucleated erythrocytes were observed in bone marrow smears of *Sec23b^ki/ko^* mice (200 cells were observed for each mouse, 2 mice for each genotype) (Fig. 3D). Therefore, the mild anemia phenotype in *Sec23b^ki/ko^* mice is distinct from the human CDAII phenotype.

**Figure 3.**
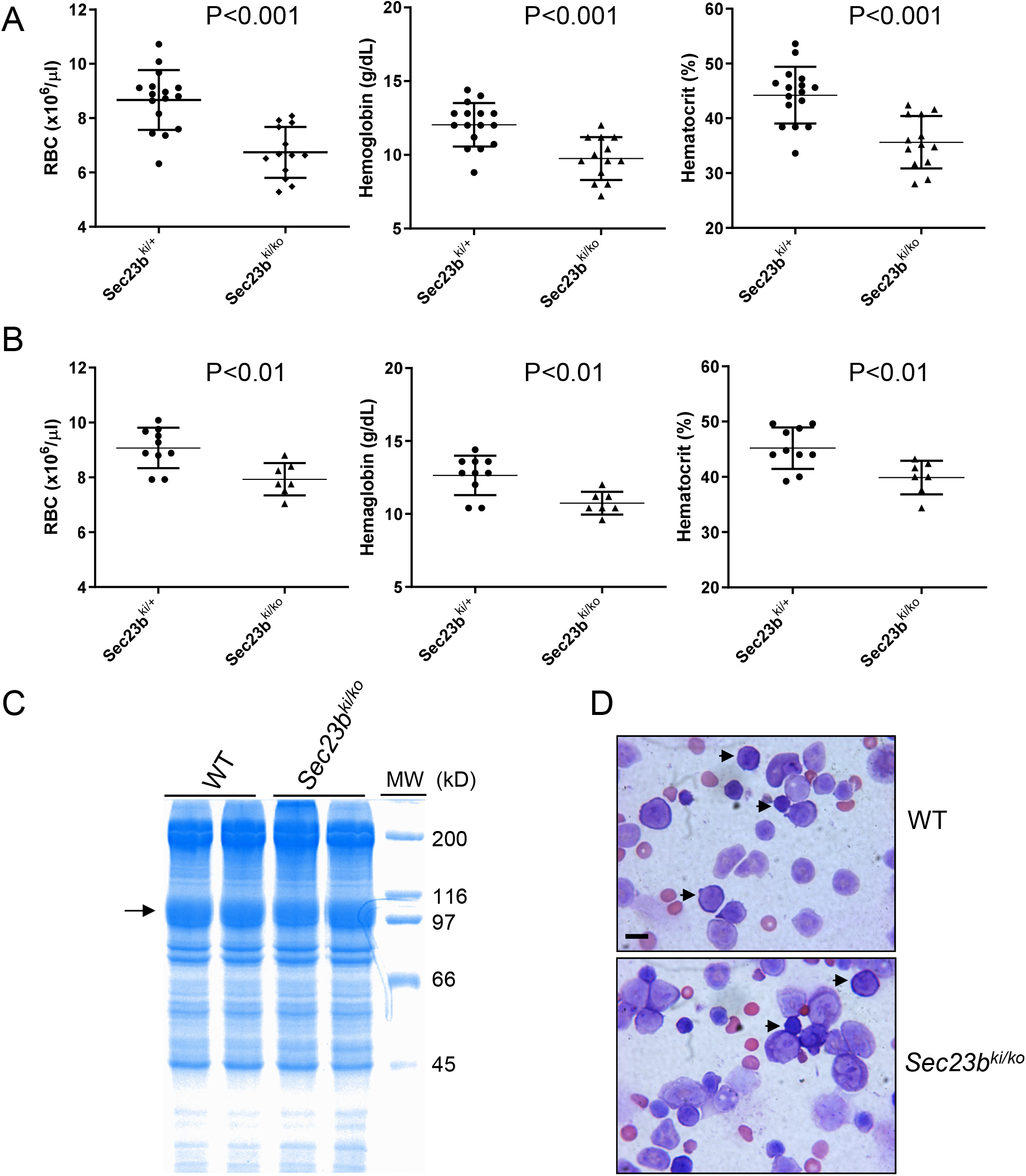
Mild to moderate anemia in *Sec23b^ki/ko^* mice without CDAII-like phenotype. A) RBC count, hemoglobin level and hematocrit from 1-month (top) and 4-month (bottom) old *Sec23b^ki/+^* and *Sec23b^ki/ko^* mice (data are mean ± SD). B) RBC ghosts were isolated from 1-month old WT (n=2) and *Sec23b^ki/ko^* mice (n=2) and separated by SDS-PAGE. Coomassie blue staining identified RBC membrane protein Band 3 (arrow). C) Wright’s stain of bone marrow smears of 4-month old WT and *Sec23b^ki/ko^* mice. Arrows indicate erythroblasts. Scale bar: 10 μm.

### Sec23b^E129K^ hemizygosity results in pancreatic insufficiency

The ratio of pancreas weight/body weight is ~40% lower for *Sec23b^ki/ko^* mice than for WT and *Sec23b^ki/ki^* mice, whereas the ratio of kidney weight/body weight is not significantly different between all groups (Fig. 4A). Histologic evaluation of pancreatic tissues of *Sec23b^ki/ko^* mice demonstrated disruption of normal lobular structure because of multifocal degeneration of exocrine epithelia cells, prominence of fat tissue and interstitial fibrosis (Fig. 4B). Masson trichrome staining detected little blue stained fibrous tissue in WT and *Sec23b^ki/ki^* mouse pancreas (Fig. 4C). However, large amounts of blue staining were readily observed in *Sec23b^ki/ko^* mouse pancreas (Fig. 4C), which further reveals exocrine cell degeneration and pancreas fibrosis. In addition, we examined the inflammatory status of the pancreas by CD45 immunohistochemistry staining, which marks total white blood cells. Compared with scant CD45 staining in WT and *Sec23b^ki/ki^* pancreas, a large number of white blood cells could be detected in *Sec23b^ki/ko^* pancreas and they were mainly distributed in the area between gland tissues where increased fibrous and fat tissue are localized (Fig. 4C). We also measured the blood pro-inflammatory cytokines TNFα, IL-1 and IL-6. While TNFα and IL-1 levels were below detection limits, the IL-6 level was significantly elevated in *Sec23b^ki/ko^* mice compared to their WT littermates (Fig. S6A). Complete deficiency in SEC23B also leads to degeneration of other professional secretory tissues (26). We compared the H&E staining of salivary glands of WT and *Sec23b^ki/ko^* mice and found no obvious defects (Fig. S6B), suggesting that pancreas is more sensitive to SEC23B deficiency.

**Figure 4.**
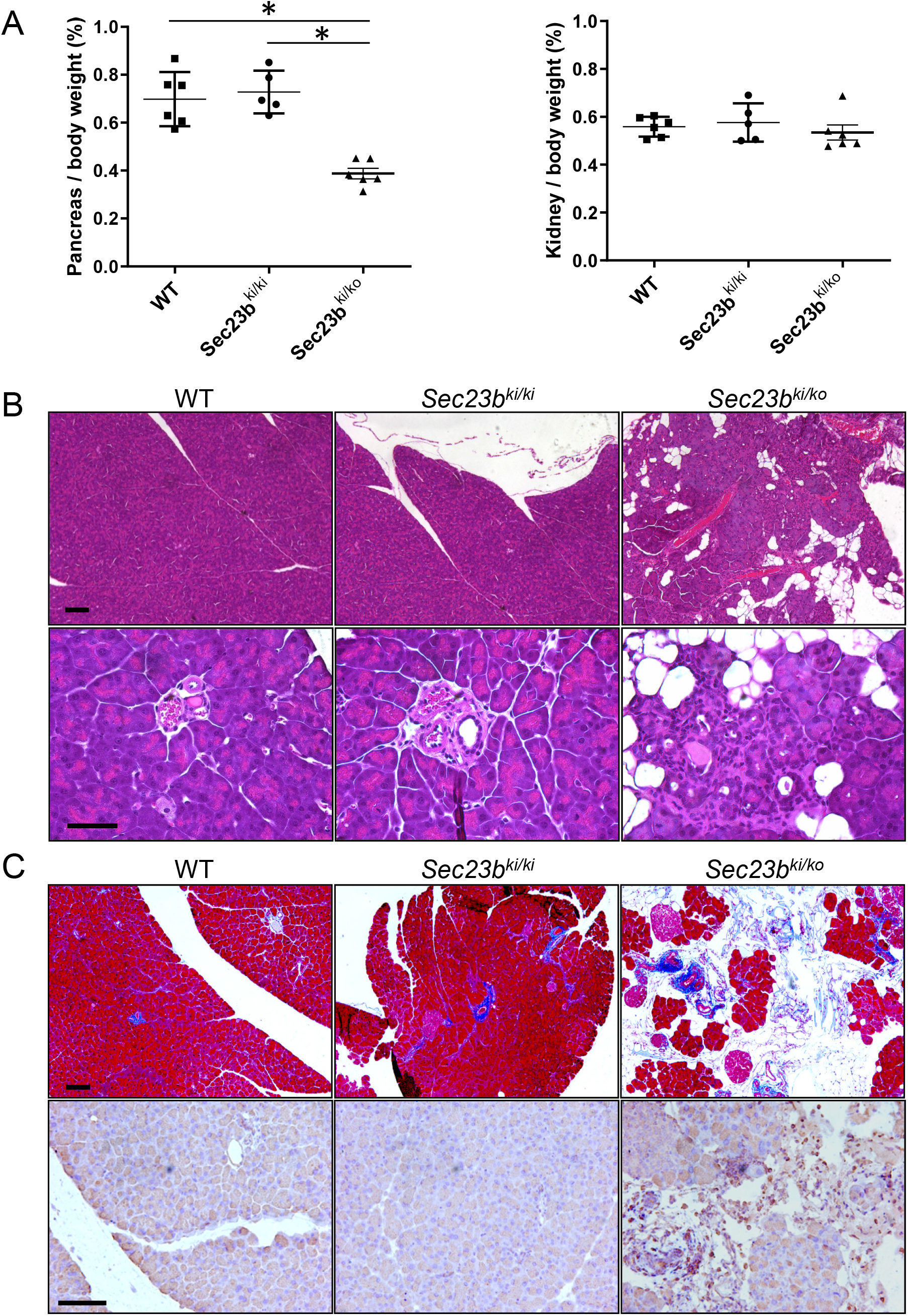
Abnormal pancreas morphology of *Sec23b^ki/ko^* mice. A) Ratios of pancreas weight/body weight and kidney weight/body weight in WT, *Sec23b^ki/ki^*, and *Sec23b^ki/ko^* mice. B) H&E staining of pancreas from WT, *Sec23b^ki/ki^* and *Sec23b^ki/ko^* mice. Scale bars: 100 μm (top) and 50 μm (bottom). C) Masson Trichrome staining of pancreas from WT, *Sec23b^ki/ki^* and *Sec23b^ki/ko^* mice. Scale bar: 100 μm. D) Immunohistochemistry staining of CD45 in pancreas from WT, *Sec23b^ki/ki^* and *Sec23b^ki/ko^* mice. Representative images from at least 3 biological replicates were shown. Scale bar: 100 μm. All mice were 2-month old.

Next, we assessed the exocrine and endocrine functions of *Sec23b^ki/ko^* mouse pancreas. To examine the exocrine function of pancreas, we collected dry or fresh stools and detected the amount of fecal protein and protease, which can indirectly assess pancreatic exocrine function (Fig. 5A). Over two fold more protein was detected in *Sec23b^ki/ko^* mouse stools than in WT and *Sec23b^ki/ki^* mouse stools (P< 0.05). Meanwhile, fecal protease level decreased over two fold in *Sec23b^ki/ko^* mouse stools compared to WT and *Sec23b^ki/ki^* mouse stools (Fig. 5A). However, serum amylase and lipase levels were not significantly different between WT, *Sec23b^ki/ko^* and *Sec23b^ki/ki^* mice (Fig. 5B). These experiments reveal significant deficiency in protein digestion in *Sec23b^ki/ko^* mouse without significant accumulation of pancreatic enzymes in blood. To assess endocrine functions of the pancreas, we measured fasting glucose and performed glucose tolerance tests (GTT) of WT and *Sec23b^ki/ko^* mice. After 6 h fasting, *Sec23b^ki/ko^* mice exhibited mild to moderate hypoglycemia in both male and female mice (Fig. 5C). However, no significant difference was observed in GTT at any time points between WT and *Sec23b^ki/ko^* mice (Fig. 5D). Immunostaining for insulin and glucagon demonstrated normal α and β cell distribution in *Sec23b^ki/ko^* mouse islets (Fig. S7).

**Figure 5.**
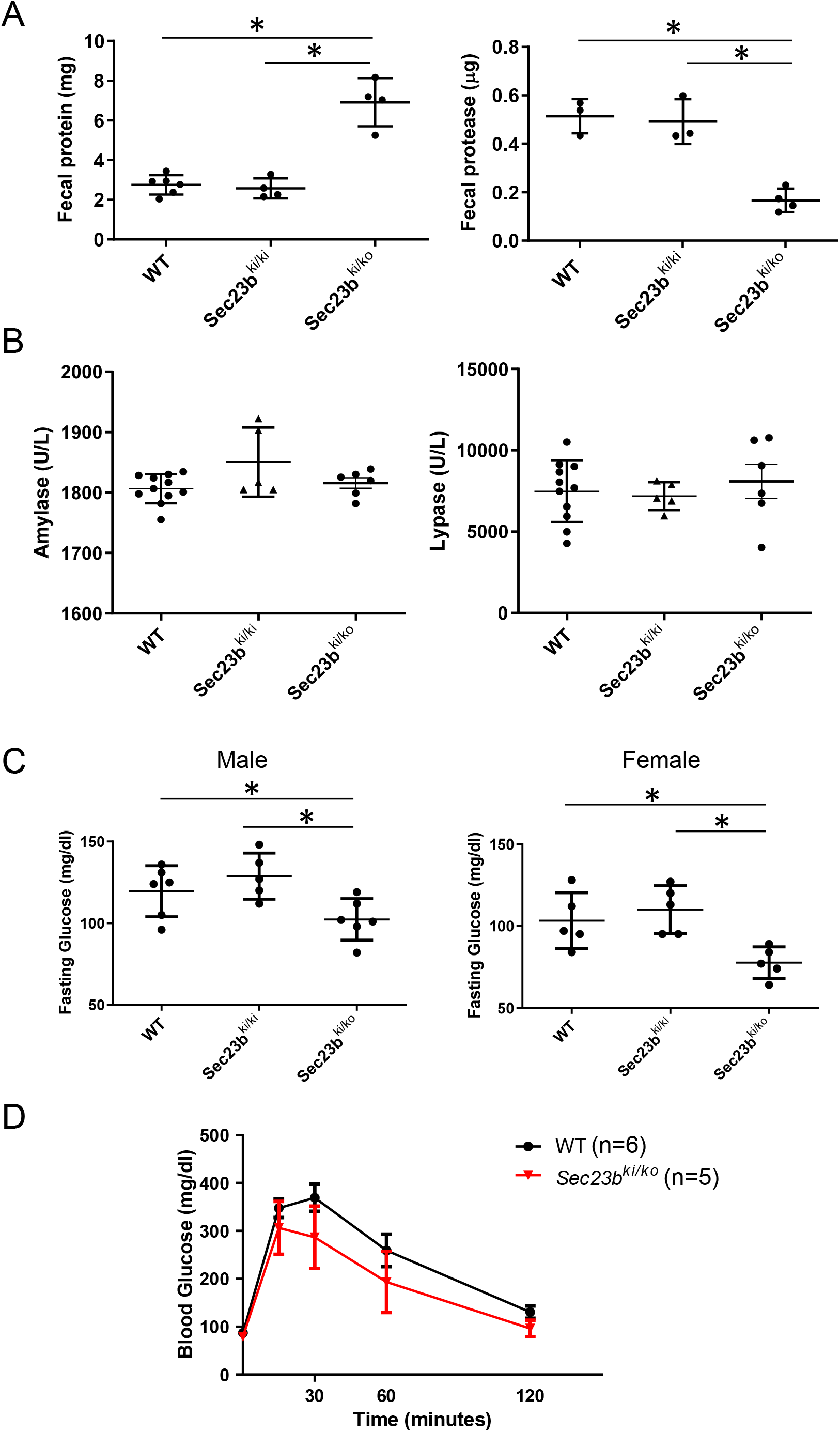
Exocrine and endocrine functions in WT, *Sec23b^ki/ki^* and *Sec23b^ki/ko^* mice. A) Fecal protein and protease levels in WT, *Sec23b^ki/ki^* and *Sec23b^ki/ko^* mice (data are mean ± SD, asterisks: P<0.05). B) Blood glucose levels after 6 hour starvation in both male and female mice of WT, *Sec23b^ki/ki^* and *Sec23b^ki/ko^* genotypes (data are mean ± SD, asterisks: P<0.05). C) Glucose tolerance test was conducted in male WT and *Sec23b^ki/ko^* mice after overnight fasting. No difference in glucose levels were found at any time points between WT and *Sec23b^ki/ko^* mice. All mice were 4-month old.

### Sec23b^E129K^ hemizygosity leads to ER stress and apoptosis of pancreatic cells

We first measured mRNA levels of genes in the ER stress pathway in WT and *Sec23b^ki/ko^* mouse pancreas. Levels of *Hspa5* (encoding GRP78), *Ddit3* (encoding CHOP) and *Trib3* (encoding TRB3) increased by 1.5-2.0 folds in *Sec23b^ki/ko^* pancreas compared with those in WT pancreas (Fig. 6A). CHOP and TRB3 are associated with apoptosis. Indeed, TUNEL staining revealed significantly increased apoptosis in *Sec23b^ki/ko^* pancreas (Fig. 6B). We next assessed pancreatic acinar cell ultrastructure using transmission electron microscopy. We observed various morphologies of acinar cells in *Sec23b^ki/ko^* mouse pancreas (Fig. 6C). There are cells with normal appearance of the ER and cells with mildly distended or disrupted ER cisternae, all with normal appearing zymogen granules (ZGs). These cells were often in proximity to cells at different stages of apoptosis. ZGs were still present in earlier stage apoptotic cells and became absent in late stage apoptotic cells (Fig. 6C). The appearances of ER in zymogen-containing cells range from indistinguishable from the WT cells to mildly distended ER with disruption of normal luminal structures (Fig. 6D). Acinar morphologies in *Sec23b^ki/ko^* pancreas are in stark contrast to previously reported *Sec23b^gt/gt^* pancreas with complete SEC23B deficiency, in which acinar cells contained severely distended ER and were devoid of ZGs (26).

**Figure 6.**
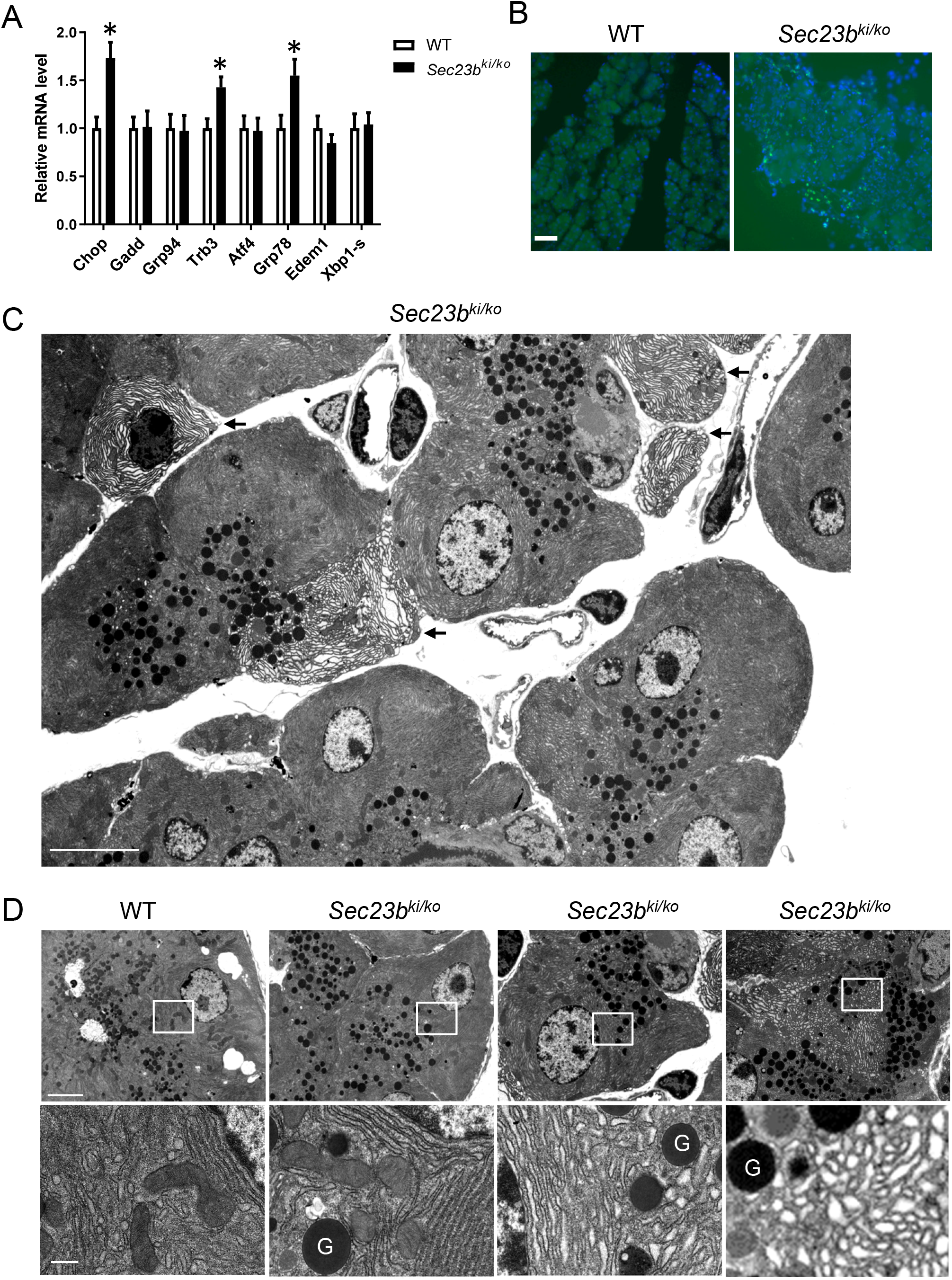
ER stress, apoptosis and ultrastructure of WT and *Sec23b^ki/ko^* mouse pancreas. A) Real-time PCR quantification was conducted for expression of selected UPR genes in pancreas of 2-month old WT (n=6) and *Sec23b^ki/ko^* (n=6) mice. Data are mean ± SD and asterisks indicate statistically significant differences between groups (P<0.05). B) Increased apoptosis in the pancreas of *Sec23b^ki/ko^* mice at 2 month of age. TUNEL staining was performed on cryosections of WT and *Sec23b^ki/ko^* pancreas. Apoptotic cells were visualized in green and nuclei are stained blue with DAPI. Scale bar: 50 μm. C) TEM image of a representative group of pancreatic acinar cells from a 2-month old *Sec23b^ki/ko^* mouse. Arrows denote apoptotic cells. Scale bar: 10 μm. D) Variant degrees of ER abnormalities of pancreatic acinar cells from a *Sec23b^ki/ko^* mouse compared to a WT mouse. G: Zymogen granule. Scale bars: 4 μm (top) and 0.5 μm (bottom). TEM images were representative of 3 mice from each genotype.

### Growth hormone (GH) insensitivity in Sec23b^ki/ko^ mice

To understand the mechanism of growth restriction in *Sec23b^ki/ko^* mice, we measured levels of two hormones important for growth, thyroxine and GH, in the serum of WT and *Sec23b^ki/ko^* mice. Thyroxine levels were similar between WT and *Sec23b^ki/ko^* mice (Fig. 7A). However, although steadily decreased as mice age, GH levels were markedly elevated in *Sec23b^ki/ko^* mice compared to WT mice (Fig. 7B). IGF-1 is a major target of GH and is mainly secreted by the liver. In contrast to GH, serum IGF-1 levels were lower in *Sec23b^ki/ko^* mice than in WT mice, and the differences were most obvious during the first 2 months of age (Fig.7C). Increased GH and decreased IGF-1 levels strongly suggest that *Sec23b^ki/ko^* mice are resistant to GH. As a major target organ of GH, liver secretes circulating IGF-1 upon GH stimulation *via* the JAK-STAT pathway. Therefore, we examined the GH receptor (GHR) and its downstream pathways including JAK-STAT, PI3K-AKT and MAPK pathways in mouse liver. As shown in Fig. 7D, GHR level decreased significantly in *Sec23b^ki/ko^* liver compared with WT liver. The intensity of phosphorylated STAT5a/b decreased to an extremely low level in *Sec23b^ki/ko^* liver (Fig. 7D), whereas the total STAT5a/b remained unchanged. On the other hand, levels of phosphorylated AKT (PI3K-AKT pathway) and phosphorylated ERK1/2 (MAPK pathway) *Sec23b^ki/ko^* liver appeared to decrease to a lesser extent (Fig. 7D). Next, we measured mRNA levels of *Ghr* and GHR-targeted genes in mouse liver by real-time RT-PCR. As shown in Fig. 7E, the mRNA levels of *Ghr* and *Igf1* consistently decreased in the liver of one, two and four-month old *Sec23b^ki/ko^* mice. The decrease was more prominent during the first 2 months. In contrast to *Ghr* and *Igf1*, the mRNA level of *Scos3*, which is a suppressor of GHR and the JAK-STAT pathway, was significantly higher in *Sec23b^ki/ko^* liver. The mRNA levels of C/EBP-β and c-FOS decreased in certain age of *Sec23b^ki/ko^* liver but not in all age groups. Therefore, different genes respond differently to the elevated GH level in *Sec23b^ki/ko^* liver.

**Figure 7.**
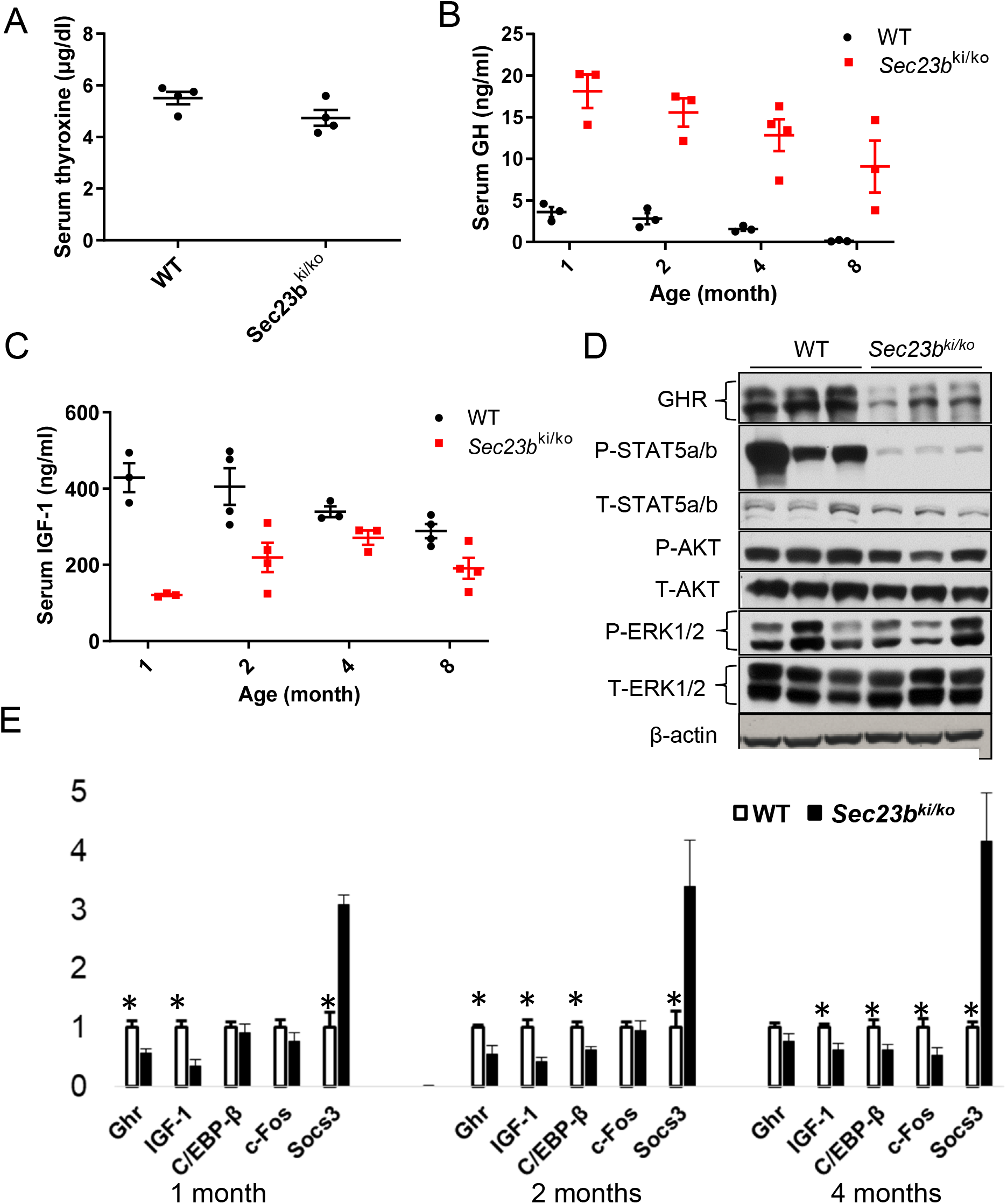
Growth hormone insensitivity in *Sec23b^ki/ko^* mice. A) Comparison of serum thyroxine levels between WT and *Sec23b^ki/ko^* mice at 2 month of age (data are mean ± SEM, P>0.05). B) Comparison of circulating GH levels between WT and *Sec23b^ki/ko^* mice from 1 to 8 months of age (data are mean ± SEM, asterisks: P<0.05). C) Comparison of circulating IGF-1 levels between WT and *Sec23b^ki/ko^* mice from 1 to 8 months of age (data are mean ± SEM, asterisks: P<0.05). D) GHR and downstream GHR signaling pathway components in liver lysates of WT and *Sec23b^ki/ko^* mice at 2 months of age. E) Real-time RT-PCR quantification of *Ghr* and select GHR target genes in WT and *Sec23b^ki/ko^* liver at 1, 2 and 4 months of age. Data are mean ± SD. Asterisks: P<0.05.

### Hepatic SEC23B deficiency is not the cause of growth restriction

To investigate whether SEC23B deficiency in liver is the main reason of growth restriction, we generated hepatocyte-specific SEC23B knockout mice. To do this, we first crossed the *Sec23b^fl/+^* mice with *Alb-cre^+^* mice that carry the Cre recombinase gene driven by the *Alb* promoter. The resulting *Sec23b^fl/+^ Alb-cre^+^* mice were then crossed with *Sec23b^fl/fl^* mice to generate *Sec23b^fl/fl^ Alb-cre^+^* hepatocyte-specific *Sec23b* knockout mice. As shown in Fig. 8A, hepatocyte-specific deletion of *Sec23b* reduced SEC23B protein to an extremely low level in *Sec23b^fl/fl^ Alb-cre^+^* mouse liver. However, unlike *Sec23b^ki/ko^* mice, we did not detect decreased GHR level in *Sec23b^fl/fl^ Alb-cre^+^* mouse liver (Fig. 8A). In contrast to *Sec23b^ki/ko^* mice, *Sec23b^fl/fl^ Alb-cre^+^* mice appeared grossly normal and did not exhibit growth restriction (Fig. 8B). Serum GH and IGF-1 levels in *Sec23b^fl/fl^ Alb-cre^+^* mice were similar to those of WT mice (Fig. 8C and 8D). Therefore, GH resistance of *Sec23b^ki/ko^* liver was not caused by hepatic SEC23B deficiency.

**Figure 8.**
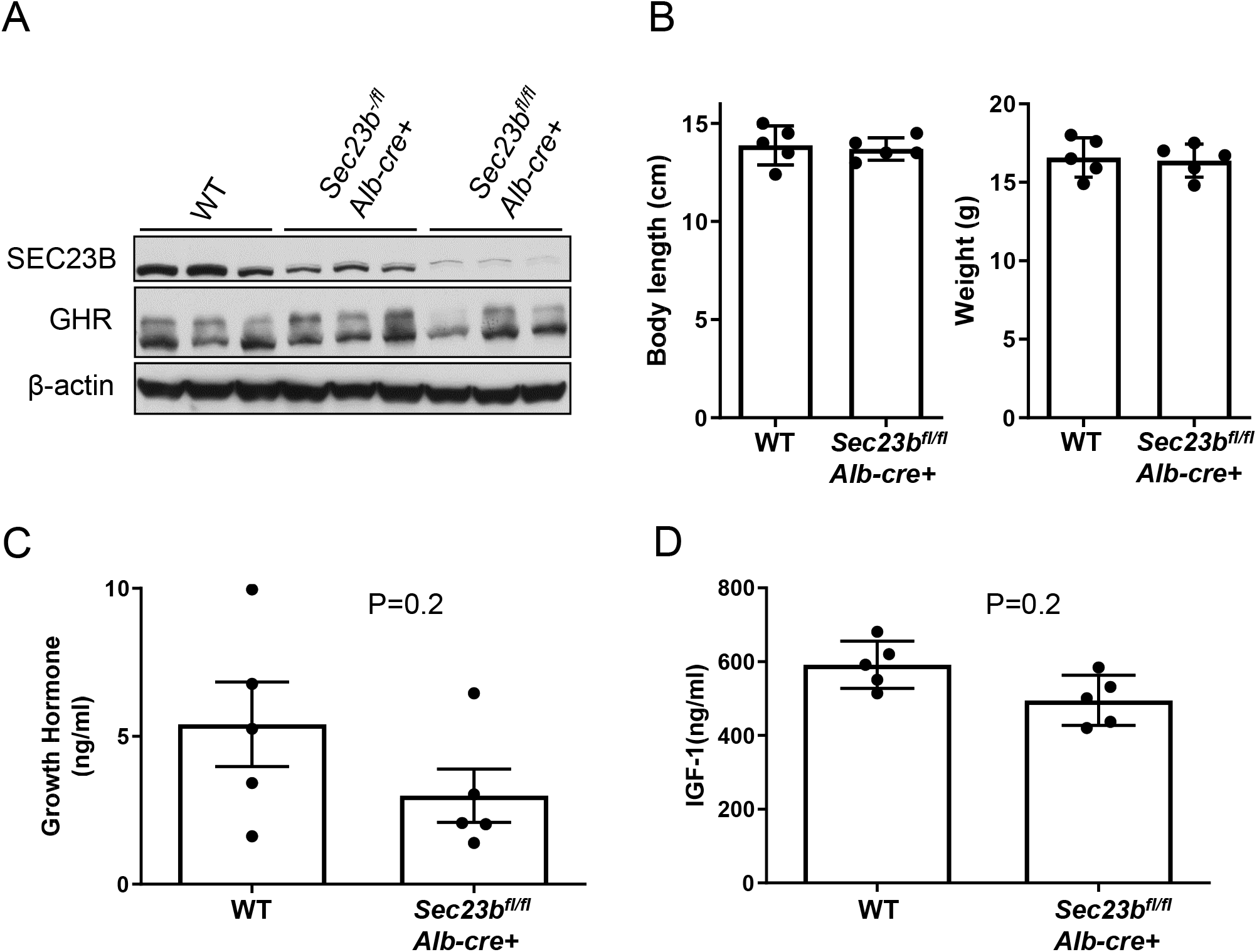
No growth restriction and growth hormone insensitivity in *Sec23b^fl/fl^ Alb-cre* mice. A) Immunoblotting analysis of SEC23B and GHR in liver lysates from 2-month old WT, *Sec23b^+/fl^ Alb-cre* (n=3) and *Sec23b^fl/fl^ Alb-cre+* (n=3) mice. B) Body weights and lengths of WT and *Sec23b^fl/fl^ Alb-cre* male mice at 5 weeks of age. C) Serum GH levels in WT and *Sec23b^fl/fl^ Alb-cre* mice. D) Serum IGF-1 levels in WT and *Sec23b^fl/fl^ Alb-cre+* mice. Both GH and IGF-1 levels were measured at 2 month of age by ELISA and showed no statistical difference (data are mean ± SEM, P>0.05).

## Discussion

Previous studies showed that mice with complete SEC23B deficiency die perinatally due to extensive degeneration of professional secretory tissues, especially pancreas (26) (29). Our new *Sec23b* knockout strain (*Sec23b^ko/ko^*) confirmed this observation. Subsequent study of mice with pancreas-specific deletion of *Sec23b* showed that the pancreatic defect is the main cause of perinatal lethality (28). Tamoxifen-inducible, pancreatic acinar cell-specific *Sec23b* deletion further demonstrated that SEC23B is required for normal function of pancreatic acinar cells in adult mice (27). However, tamoxifen-inducible, pancreatic acinar cell-specific *Sec23b* deletion is only suitable for short-term study of the impact of SEC23B on the function of the pancreas. We report the characterization of mice with homozygous and hemizygous E109K mutation of *Sec23b*. The human counterpart of this mutation is the most common missense mutation identified in CDAII patients. Mice with homozygous SEC23B^E109K^ mutant alleles (*Sec23b*^ki/ki^) are grossly normal, suggesting that SEC23B carrying the E109K mutation retains partial function. In addition, the SEC23A level is significantly elevated in *Sec23b*^ki/ki^ mice, largely compensating for the loss of SEC23B function. Further reducing the gene dosage dramatically altered the survivability and phenotype, as postnatal death and pancreatic insufficiency are associated with hemizygous SEC23B^E109K^ mutation, but not with homozygous SEC23B^E109K^ mutation. Surviving *Sec23b^ki/ko^* mice also exhibit severe growth restriction, particularly early in life. Since the number of neonatal *Sec23b^ki/ko^* pups is not significantly different from the expected number, and the weight and body length of *Sec23b^ki/ko^* neonates are normal, the E109K hemizygosity is compatible with embryonic development. Thus, the growth restriction is the result of failure to thrive after birth. The first three weeks of life are most critical, as half of *Sec23b^ki/ko^* mice died during this period. Body weight and length differences between WT and *Sec23b^ki/ko^* mice narrowed as mice age, coinciding with the decrease of SEC23B level in WT pancreas, suggesting relatively decreased reliance on SEC23B over time. In humans, phenotype severity also correlates with the residual SEC23B activity, as more severe anemic phenotype was observed in CDAII patients with a missense allele and a null allele (23).

In our study, hemizygous SEC23B^E109K^ mutation leads to chronic changes in the pancreas which have not been found in other SEC23B deficient mice, including degeneration and inflammation of the pancreatic tissue, increased fat deposition, interstitial fibrosis of the pancreas accompanied by significant infiltration of white blood cells. These changes in *Sec23b^ki/ko^* pancreas meet the criteria of morphologic changes in chronic pancreatitis. Mice with tamoxifen-inducible, acinar cell-specific SEC23B deficiency exhibit pancreatic cell loss, but without an effect on viability of the mice or significant evidence of inflammation (27). Besides histologic changes in the pancreas, chronic pancreatitis also leads to exocrine and endocrine defects in human. One of the major phenotypes of exocrine deficiency is malabsorption of lipid and protein, which can result in malnutrition in humans. In our study, deficiency in protein absorption was indirectly detected in *Sec23b^ki/ko^* mice, suggesting specific disruption of protease transport, such as chymotrypsinogen and carboxypeptidase, in pancreatic acinar cells. There is evidence that different enzymes move through the early secretory pathway to ZGs at different rates and thus may be differently affected by SEC23B deficiency (31). We did not observe increases in amylase and lipase levels in *Sec23b^ki/ko^* mice. In human chronic pancreatitis, blood amylase and lipase levels are often not elevated or only slightly elevated, except during acute attacks.

In contrast to all the reported SEC23B deficient mice so far, *Sec23b^ki/ko^* mice did exhibit a mild to moderate anemia phenotype. However, this anemia is not accompanied by other characteristic features of CDAII, including increased binucleated erythrocytes and hypoglycosylated band 3 in mature red blood cell membranes. Lack of CDAII phenotype is most likely due to SEC23A as the dominant paralog expressed in the hematopoietic cells (26) (29). The anemia phenotype is most likely caused by malnutrition as a result of pancreatic insufficiency. Similar to previous studies, no major disruption in islet structure was observed in *Sec23b^ki/ko^* mice. Interestingly, we found hypoglycemia in *Sec23b^ki/ko^* mice after fasting, while hyperglycemia is usually observed in chronic pancreatitis patients. At the same time, there was no significant difference in GTT tests, suggesting no severe destruction of islet β cell functions. Low storage of glycogen caused by malnutrition could explain the fasting hypoglycemia in *Sec23b^ki/ko^* mice. Thus, SEC23B appears to be more important for exocrine than endocrine functions of the mouse pancreas.

Blockage of this secretory pathway can result in accumulation of exocrine pancreatic enzymes in the ER which further induces ER stress. In both humans and mice, ER stress has been found in association with pancreatitis (32–35). In humans, variants in procarboxypepidase A1 and mutations in chymotrypsinogen C were associated with both chronic pancreatitis and ER stress (32, 35). In addition, alterations in the activity of trypsinogen cause hereditary pancreatitis resulting from ER stress, such as mutations in the *PRSS1* and *SPINK1* genes (33, 36). In our study, we detected mild levels of ER stress and increased apoptosis in *Sec23b^ki/ko^* pancreas. Acinar cells with different levels of ER distension and varying numbers of ZGs were observed by TEM, including acinar cells at different stages of apoptosis. This is in contrast to severe ER distension and complete lack of ZGs in total SEC23B deficient mice (26). This observation suggests that acinar cells with *Sec23b*^E129K^ hemizygosity can function for a period of time before succumbing to apoptosis induced by continued ER stress, and may explain the relatively mild pancreas degeneration. Therefore, ER stress caused by SEC23B deficiency may trigger chronic pancreatitis in *Sec23b^ki/ko^* mice.

To investigate the underlying reasons accounting for the growth restriction in *Sec23b^ki/ko^* mice, we unexpectedly found increased GH and decreased IGF-1 levels in *Sec23b^ki/ko^* mice, which strongly suggested GH insensitivity in these mice. GH plays a pivotal role in linear growth by binding to its widely expressed transmembrane receptor GHR, which subsequently activates downstream pathways and modulates the expression of its target genes. We further confirmed GH resistance by demonstrating impaired GHR signaling in *Sec23b^ki/ko^* liver. Several conditions have been found to be related to GH resistance, such as loss of function mutations in the GH gene, undernutrition, imbalance of the endocrine system and inflammation (37, 38). In humans, a disease called Laron syndrome is caused by defects in GHR (39, 40). Laron syndrome patients have similar phenotype to *Sec23b^ki/ko^* mice in growth restriction, increased GH and decreased serum IGF-1 levels. Considering the function of SEC23B in intracellular protein transport, we asked whether SEC23B deficiency results in a defect in transporting GHR. Therefore, we generated hepatic *Sec23b* conditional knockout mice. Liver-specific deletion of GHR resulted in more than 90% suppression of serum IGF-1 and an over 3-fold increase in GH level (41). However, we did not observe significant differences in IGF-1 and GH levels between WT and *Sec23b*^-/-^ *Alb-cre+* mice, suggesting normal GHR transport in SEC23B deficient hepatocytes.

GH insensitivity can occur in inflammatory states in both clinical and experimental settings (42–46). Previous studies demonstrated that exogenous pro-inflammatory cytokines such as IL-6, TNF-α and IL-1β inhibit GH signaling *in vitro* and *in vivo* (47–49). As discussed above, chronic pancreatitis in *Sec23b^ki/ko^* mice might be the major cause of GH resistance by generating a whole body chronic inflammatory state. Consistent with this hypothesis, we observed elevated IL-6 levels in *Sec23b^ki/ko^* mice. In addition, we found increased *Socs3* mRNA level in *Sec23b^ki/ko^* liver. SOCS3 is a major mediator of inflammation-induced GH resistance in liver (42, 48, 50). Upon inflammatory cytokine stimulation, IL-6 in particular, SOCS3 down regulates the JAK2-STAT5 pathway in several ways such as inhibiting *Ghr* transcriptional activity and inhibiting GH activation of the signal transducer STAT5b by binding with GHR’s membrane-proximal tyrosine residues (42, 51). Consistent with the well-known effect of SOCS3 on JAK2/STAT5, we detected a dramatic decrease in phosphorylated STAT5 levels in *Sec23b^ki/ko^* liver. It is also well known that pro-inflammatory cytokines are able to reduce GHR mRNA and protein levels both *in vivo* and *in vitro* (43, 47, 52, 53), consistent with the decreased GHR level observed in *Sec23b^ki/ko^* liver. Therefore, pro-inflammatory cytokines triggered by chronic pancreatitis could be one of the main reasons for the GH resistance observed in our mice.

## Methods

### Generation of Sec23b conditional knockout (KO) mice

The *Sec23b* KO mice were produced from a vector that was designed to delete exons 5 and 6 of *Sec23b* in C57BL/6 ES cells. The 5’ homology arm (~4.6 kb containing exons 3 and 4), the 3’ homology arm (~5.0 kb containing exons 5-9) and the center piece (~1.0 kb containing exons 5 and 6) were amplified by PCR using C57BL/6 bacterial artificial chromosome (BAC) DNA as a template and cloned into the targeting vector LoxFtNwCD. The targeting construct was linearized using NotI and electroporated into C57BL/6 ES cells. The targeted allele was identified by Southern blot. Mice with germline transmission of the targeting allele (*Sec23b^fl/+^* mice) were continually crossed with C57BL/6J mice. Global KO mice were generated by crossing *Sec23b^fl/+^* mice with EIIa-cre mice to delete exons 5 and 6 and the neo cassette flanked by the LoxP sequences. Hepatocyte-specific mice were generated by crossing *Sec23b^fl/+^* mice with *Alb-cre^+^* mice.

### Generation of Sec23b^E109K^ knockin (KI) mice

To produce the *Sec23b*^E109K^ KI mice, 5’ homology arm (~3.2 kb) and 3’ homology arm (~7.2 kb) were cloned by PCR using C57BL/6 BAC DNA as a template. The E109K mutation was introduced into exon 4 (in the 5’ arm) by site-directed mutagenesis. In the targeting vector, the *Neo* cassette was flanked by LoxP sites. Homologous recombination was performed on C57BL/6 ES cells to identify the targeted allele. The targeted allele was identified by Southern blot. The *Neo* cassette was removed by transfection of the targeted clones with a *Cre*-expressing plasmid. Mice with germline transmission of the targeting allele were continually crossed with C57BL/6J mice.

### Mouse genotyping

Genotyping of *Sec23b* KI mice was carried out by a two-primer PCR assay of genomic DNA prepared from tail clippings of pups (primers: CCF19 Neo F and CCF19 Neo R). A three-primer PCR assay was used to genotype *Sec23b* KO mice (primers: CCF13 DelR, CCF13 LoxPR and CCF13 LoxPF). Primer sequences are listed in Supplemental Table 1.

### Animal procedures

Blood glucose levels were measured using the Contour Blood Glucose Monitoring System (Bayer, Leverkusen, Germany). The complete blood count was determined in an Advia120 whole blood analyzer (Bayer, Leverkusen, Germany). BM was flushed from femurs and tibias of each mouse with Hanks’ balanced salt solution. Bone marrow cytospin smear was stained by Wright’s staining. Glucose tolerance tests (GTT) were performed on 4 months old mice after overnight fasting. The mice were injected intraperitoneally with a D-(+)-glucose bolus (2 g/kg of body weight). A small incision was made at the distal end of the tail vein, and blood glucose levels were measured before and at 15, 30, 60, 120 min after glucose injection. Animals were re-fed immediately after the test.

### Measurement of total protein and proteases in mouse feces

Total proteases in fresh mouse feces were measured as previously reported (54). To measure total fecal protein, 200 mg of dried feces were homogenized in 3 ml water and incubated at 4 °C overnight. Samples were then centrifuged at 15,000 g at 4 °C for 20 min. The supernatant was collected and stored at 4°C. Next, 2 ml of 0.1 N NaOH was added to the pellet and the mixture was gently rocked at room temperature for 1 h before centrifugation at 15,000g at 4°C for 20 min. This supernatant was combined with the first supernatant. 25% trichloroacetic acid (TCA) was added to the combined supernatant with the ratio of 2.5:1. The sample was kept in ice for 30 minutes and centrifuged at 15,000g for 15 mins at 4°C. The supernatant was discarded and the pellet was rinsed with cold 10% TCA followed by 5% TCA. The pellet was solubilized with 1 N NaOH. Protein concentrations were determined by the Bradford assay.

### RBC ghost preparation

One hundred microliters of peripheral blood was centrifuged at 3,000 rpm in a microfuge. The pellet was washed twice with PBS and then lysed by suspension in ghost lysis buffer (5 mM Na_2_PO_4_, 1.3 mM EDTA; pH 7.6) containing protease inhibitors. Lysates were centrifuged at 16,000 g, and the pellets containing the RBC membrane fraction were collected and washed five times in ghost lysis buffer.

RBC ghosts were analyzed by SDS-PAGE and proteins were visualized by Commassie blue staining.

### Antibodies

Rabbit polyclonal anti-SEC23A and anti-SEC23B antibodies were purchased from MilliporeSigma (Burlington, MA, ABC424 and ABC460). Rabbit polyclonal antibody against both SEC23A and SEC23B (anti-COPII, PA1-069A) was purchased from ThermoFisher (Waltham, MA). Anti-GHR antibody was purchased from Santa Cruz (Dallas, TX, sc-137185), and anti-β-actin was from Sigma Aldridge (St. Louis, MO, A5441). P-STAT5a/b, T-STAT5 a/b, P/T-AKT and P/T ERK ½ were all purchased from Cell Signaling (Danvers, MA).

### Histological, Masson trichrome stain and TUNEL staining

Tissues were fixed in 10% neutral buffered formalin solution (Fisher Scientific, Waltham, MA), embedded in paraffin, and cut into 5-μm–thick sections before hematoxylin and eosin (H&E) staining. Images were visualized and captured with a Zeiss Axioplan2 imaging microscope. Masson trichrome staining kit was purchased from Sigma Aldridge (St. Louis, MO) and was performed according to the provided protocol. In cell death experiments, apoptotic cells in pancreas frozen sections were detected by the TUNEL (TdT mediated dUTP nick end labeling) assay using a fluorescein-based detection kit (*in situ* death detection kit, Roche, Indianapolis, IN) according to manufacturer’s instructions. Sections were then examined under an inverted fluorescence microscope (Leica, Wetzlar, Germany).

### Electron microscopy

Small pieces of pancreas were fixed in 2.5% glutaraldehyde and 4% formaldehyde for 24 h, followed by post fixation in 1% osmium tetroxide for 1 h. After *en bloc* staining and dehydration in a graded ethanol series, samples embedded in eponate 12 medium (Ted Pella, Redding, CA). Ultrathin sections (85 nm) were doubly stained with 2% uranyl acetate and 1% lead citrate, and then observed using a PhilpsCM12 transmission electron microscope at an accelerating voltage of 60 kV by a person blind to the genotypes.

### RNA preparation and real-time reverse transcriptase (RT)-PCR

Total RNA was extracted using the Trizol reagent (ThermoFisher, Waltham, MA) followed by purification using the RNA Mini kit (ThermoFisher, Waltham, MA). RNA quantity and purity were determined by a Nanodrop Spectrophotometer. The total RNA (1 μg) from each sample was reverse transcribed into cDNA using the iScript cDNA select Synthesis Kit (Bio-Rad, Hercules, CA), according to the manufacturer’s instructions. SYBR green based quantitative PCR reactions were performed in a Bio-Rad CFX96 Real Time PCR Detection System. Reaction specificity was determined by product melting curves. Relative gene expression was calculated by the 2^-ΔΔCt^ method using *Gapdh* as a reference gene. Primer sequences are listed in Supplemental Table 1.

### Immunoblotting and Immunoprecipitation

Immunoblotting and immunoprecipitation protocols were described recently (55). Images of chemiluminescent blot signals from horseradish peroxide-conjugated secondary antibodies (Bio-Rad, Hercules, CA) were acquired by exposure to X-ray films or through the Amersham Imager 600 (GE Healthcare, Little Chalfont, U.K.) and quantified using the ImageJ software.

### Serum growth hormone (GH), IGF-1, thyroine, amylase and lipase levels

ELISA was performed to detect serum GH (rat/mouse growth hormone ELISA kit, Merck KGaA, Dermstadt, Germany), IGF-1 (mouse/rat IGF-1 ELISA kit, R&D system, Minneapolis, MN) and total thyroxine (total thyroxine ELISA, ALPCO, Salem, NH) according to protocols of the respective ELISA kits. Serum amylase and lipase activities were measured using kits from BioAssay Systems (ECAM-100 and DLPS-100, Hayward, CA) according to manufacturer’s instructions.

### Statistical analysis

All data are presented as mean ± standard deviation (SD) or standard error of the mean (SEM), as indicated in figure legends. Two-tailed Student’s *t*-test was used for comparison of continuous variables and Chi-square test was used for comparison of binary variables. All *P* values were two-sided and *P* <0.05 was considered statistically significant.

### Study approval

Animal experimental protocols were approved by the institutional animal care and use committee of Cleveland Clinic Lerner Research Institute.

## Acknowledgement

This work was supported in part by grants from the National Institutes of Health (RO1 HL094505 to B.Z. and R01 HL148333 to R.K.), and from the National Natural Science Foundation of China (81860430 to MZ). B.Z. was also supported by a grant from the Lisa Dean Moseley Foundation. R.K. was also supported by the University of Michigan Rogel Cancer Center P30CA046592 grant. We thank Yin Mei at the LRI Imaging Core for her expert support in electron microscopy.

## Authorship

Contributions: WW, MZ and BZ conceived and designed the study. WW and ZL performed most experiments and generated data. CZ and MZ performed part of experiments. WW, ZL, RK, MZ and BZ analyzed data. WW and BZ wrote the manuscript. All authors were involved in writing and editing of the manuscript.

