## Supplemental file for "A common SEC23B missense mutation in congenital dyserythropoietic anemia type II leads to growth restriction and symptoms of chronic pancreatitis in mice"

### Supplemental methods

*Cell culture.* The Nthy-ori 3-1 human thyroid follicular epithelium cell line (catalog no. EC90011609, lot no. 09C008, passage no. 16, purchased in 2014 from Sigma) was cultured in RPMI-1640 supplemented with 2 mM glutamine and 10% FBS. All cell lines were maintained at 37 ºC and 5% CO2 culture conditions and tested negative upon routine mycoplasma testing with the MycoAlert Mycoplasma Detection Kit (Lonza) at the C.E. lab (luminescence ratios < 0.9).

*Plasmids, mutagenesis, cell-Line transfection and transduction.* A retroviral plasmid (pMSCV) expressing C-terminally GFP-tagged wild-type SEC23B was previously reported (1). Wild-type plasmids were mutagenized for the missense mutations of E109K and R14W with the QuikChange II Site-Directed Mutagenesis Kit (Agilent Technologies). All expression constructs were validated by Sanger sequencing prior to transduction. To generate stable cell lines, we kept retrovirally transduced cells under 1 mg/ml puromycin selection for >30 days prior to downstream experiments.

*Immunofluorescence staining.* For Nthy-ori 3-1 immunofluorescence staining (1), cells were seeded on coverslips and were fixed with 4% paraformaldehyde for 10 min at room temperature. Coverslips were blocked in 5% BSA, permeabilized in 0.3% Triton X-100, and then incubated overnight with the primary antibodies. Fluorescent secondary antibodies conjugated with Alexa 488 or Alexa 594 (ThermoFisher) were used for signal detection. Cellular nuclei were counterstained with DAPI. Coverslips were visualized and images were obtained with a TCS SP5 confocal microscope (Leica). For immunofluorescence analysis of insulin and glucagon (2), pancreatic tissues were fixed in 4% paraformaldehyde, washed and incubated in 30% sucrose, before cryo-embedding. Sagittal, transverse and coronal 5-μm-thick sections or fixed pancreas were blocked in 5% BSA, permeabilized in 0.3% Triton X-100, and then incubated overnight with the primary antibodies. Fluorescent secondary antibodies conjugated with Alexa 488 or Alexa 594 was used for signal detection. Cellular nuclei were counterstained with 4,6-diamidino-2-phenylindole (Dapi). Sections were then examined under an inverted fluorescence microscope (Leica).

*Characterization of E18.5 Sec23b$^{ko/ko}$ embryos*. Time mating of *Sec23b*$^{ko/+}$ X *Sec23b*$^{ko/+}$ cross, dissection of pancreas from E18.5 embryos, and H&E staining of pancreas cryosections were performed as previously reported (2).

*Measurements of pro-inflammatory cytokines*. Serum TNFα, IL-1 and IL-6 levels were measured using ELISA kits (R&D Systems) according to manufacturer instructions.

Table S1. Primers used in the study.

| Primer name | Sequence (5' to 3') | Use |
|---|---|---|
| CCF19 NeoF | GGAGCACCAATCACTTTGAGCCC | KI mouse genotyping |
| CCF19 NeoR | AAGGGAGATCATGGGAAGGGTGG | KI mouse genotyping |
| CCF13 LoxPF | GATAGACTCTGGGTCCTCATCATTGGA | KO mouse genotyping |
| CCF13 LoxPR | GACATAACCAGCGAGCACAGAGAGAC | KO mouse genotyping |
| CCF13 delR | CAACAGCAATGGACAAAGCAACAC | KO mouse genotyping |
| Gapdh-S | AGGTCGGTGTGAACGGATTTG | qRT-PCR for Gapdh gene |
| Gapdh-AS | TGTAGACCATGTAGTTGAGGTCA | qRT-PCR for Gapdh gene |
| Xbp1-sF | GAGTCCGCAGCAGGTG | qRT-PCR for spliced Xbp1 gene |
| Xbp1-sR | GTGTCAGAGTCCATGGGA | qRT-PCR for spliced Xbp1 gene |
| Atf4-F | ATGGCCGGCTATGGATGAT | qRT-PCR for Atf4 gene |
| Atf4-R | CGAAGTCAAACTCTTTCAGATCCATT | qRT-PCR for Atf4 gene |
| Edem1-F | GCAATGAAGGAGAAGGAGACCC | qRT-PCR for Edem1 gene |
| Edem1-R | TAGAAGGCGTGTAGGCAGATGG | qRT-PCR for Edem1 gene |
| Gadd34-F | CCCGAGATTCCTCTAAAAGC | qRT-PCR for Ppp1r15a gene |
| Gadd34-R | CCAGACAGCAAGGAAATGG | qRT-PCR for Ppp1r15a gene |
| Grp78-F | CATGGTTCTCACTAAAATGAAAGG | qRT-PCR for Hspa5 gene |
| Grp78-R | GCTGGTACAGTAACAACTG | qRT-PCR for Hspa5 gene |
| Grp94-F | TCGTCAGAGCTGATGATGAAGT | qRT-PCR for Hsp90b1 gene |
| Grp94-R | GCGTTTAACCCATCCAACTGAAT | qRT-PCR for Hsp90b1 gene |
| Chop-F | CTGGAAGCCTGGTATGAGGAT | qRT-PCR for Ddit3 gene |
| Chop-R | CAGGGTCAAGAGTAGTGAAGGT | qRT-PCR for Ddit3 gene |
| Trb3-F | TCTCCTCCGCAAGGAACCT | qRT-PCR for Trib3 gene |
| Trb3-R | TCTCAACCAGGGATGCAAGAG | qRT-PCR for Trib3 gene |
| Ghr-F | ACAGTGCCTACTTTTGTGAGTC | qRT-PCR for Ghr gene |
| Ghr-R | GTAGTGGTAAGGCTTTCTGTGG | qRT-PCR for Ghr gene |
| Igf1-F | TCAGACAGGCATTGTGGATGAG | qRT-PCR for Igf1 gene |
| Igf1-R | GGACGGGGACTTCTGAGTCTT | qRT-PCR for Igf1 gene |
| Cebpb-F | ACCGGGTTTCGGGACTTGA | qRT-PCR for Cebpb gene |
| Cebpb-R | GTTGCGTCAGTCCCGTGTCCA | qRT-PCR for Cebpb gene |
| c-Fos-F | CGGGTTTCAACGCCGACTA | qRT-PCR for Fos gene |
| c-Fos-R | TTGGCACTAGAGACGGACAGA | qRT-PCR for Fos gene |
| Socs3-F | ATGGTCACCCACAGCAAGTTT | qRT-PCR for Socs3 gene |
| Socs3-R | TCCAGTAGAATCCGCTCTCCT | qRT-PCR for Socs3 gene |

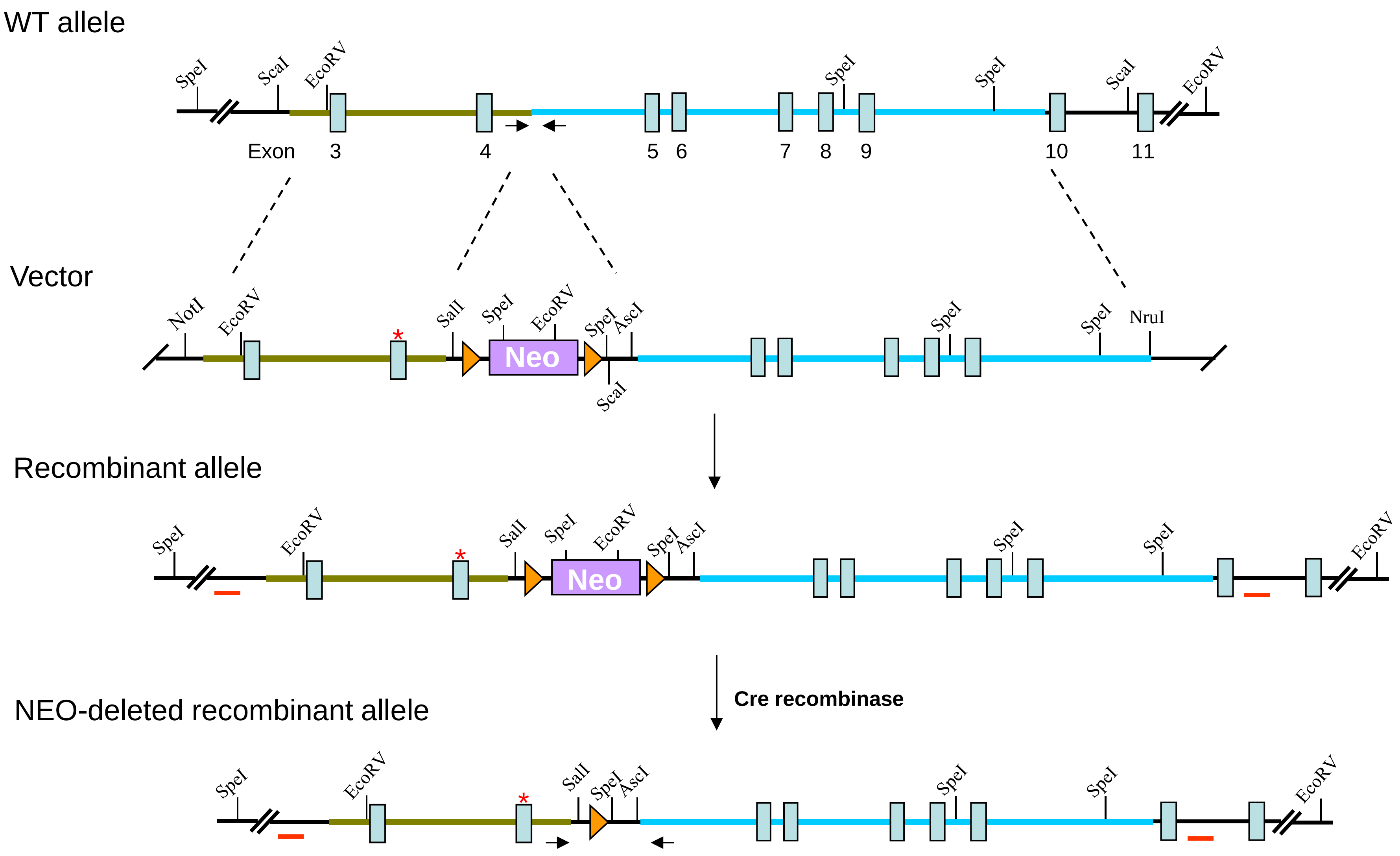


Figure S1. Detailed diagram of *Sec23b*$^{E109K}$ knockin mouse generation. Asterisks indicate the location of E109K mutation. Arrows denote locations of genotyping primers. Short red bars denote locations of Southern blot probes.

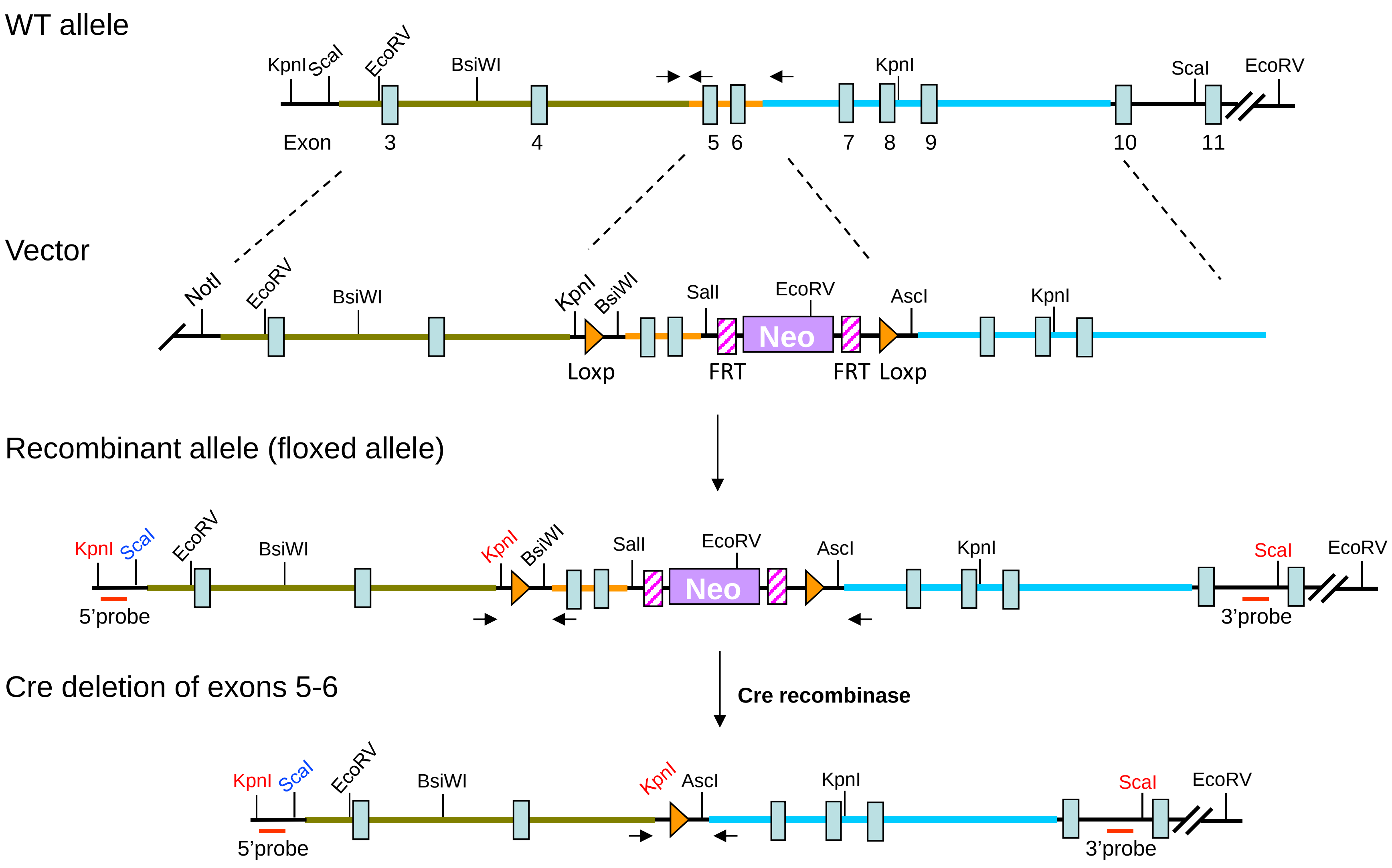


Figure S2. Detailed diagram of *Sec23b* conditional knockout mouse generation. Arrows denote locations of genotyping primers. Short red bars denote locations of Southern blot probes.

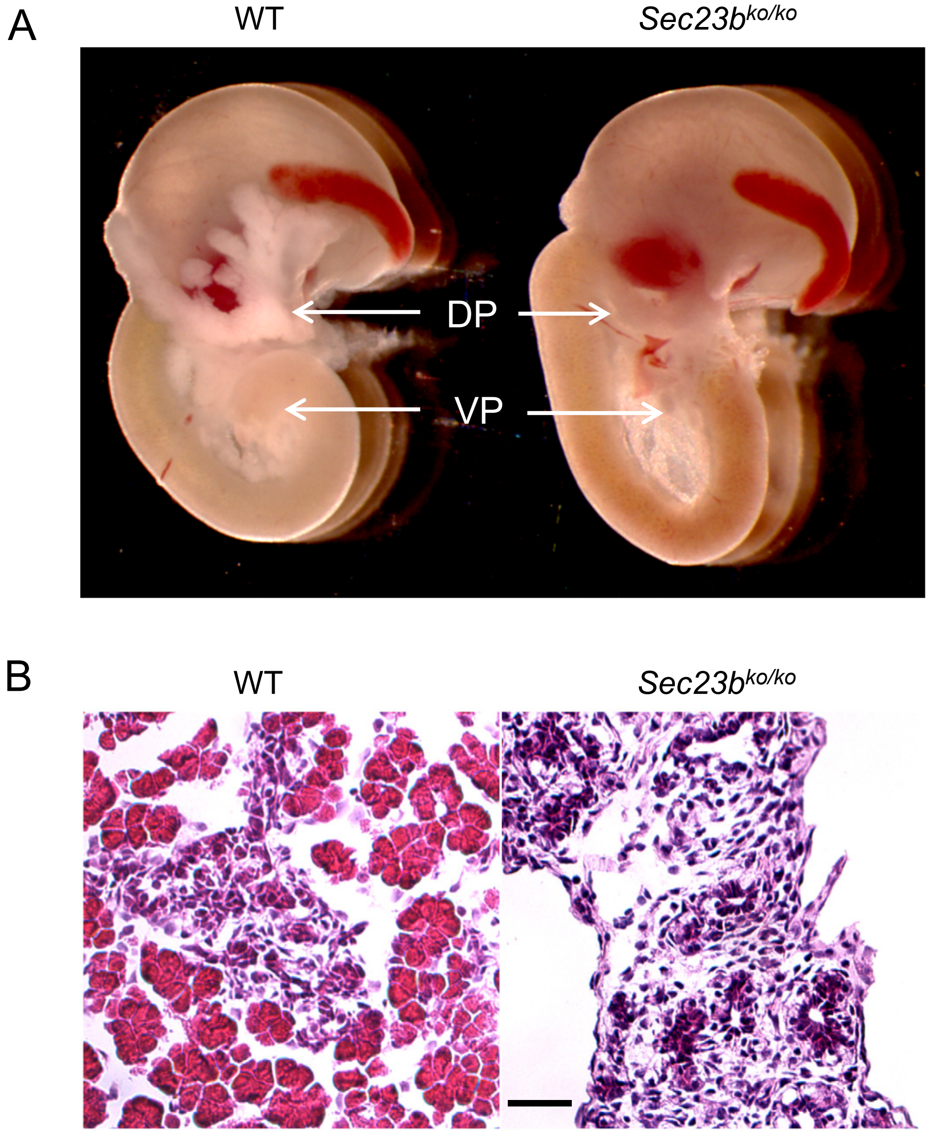


Fig. S3. Pancreas defect in E18.5 *Sec23b*$^{ko/ko}$ embryos. (A) Pancreas tissues dissected from E18.5 embryos are smaller and less opaque than from WT embryos. DP, dorsal pancreas; VP; ventral pancreas. (B) H&E staining of WT and *Sec23b*$^{ko/ko}$ pancreas cryosections demonstrates extensive destruction of pancreatic parenchyma. Scale bar: 50 µm.

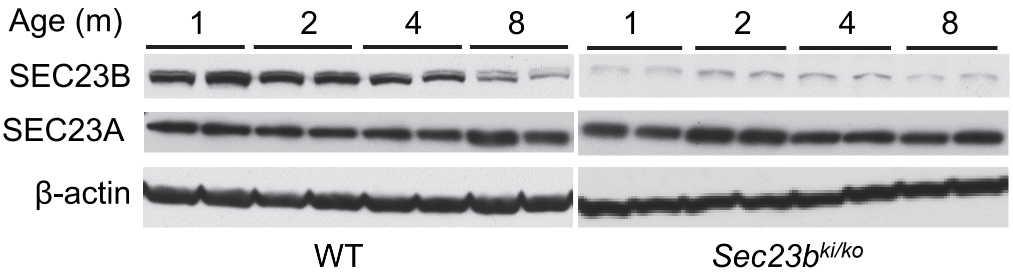


Fig. S4. Immunoblotting of SEC23B, SEC23A and β-actin in pancreas from WT and *Sec23b*$^{ki/ko}$ mice of the indicated ages. Two mice of each genotype were analyzed for each time point.

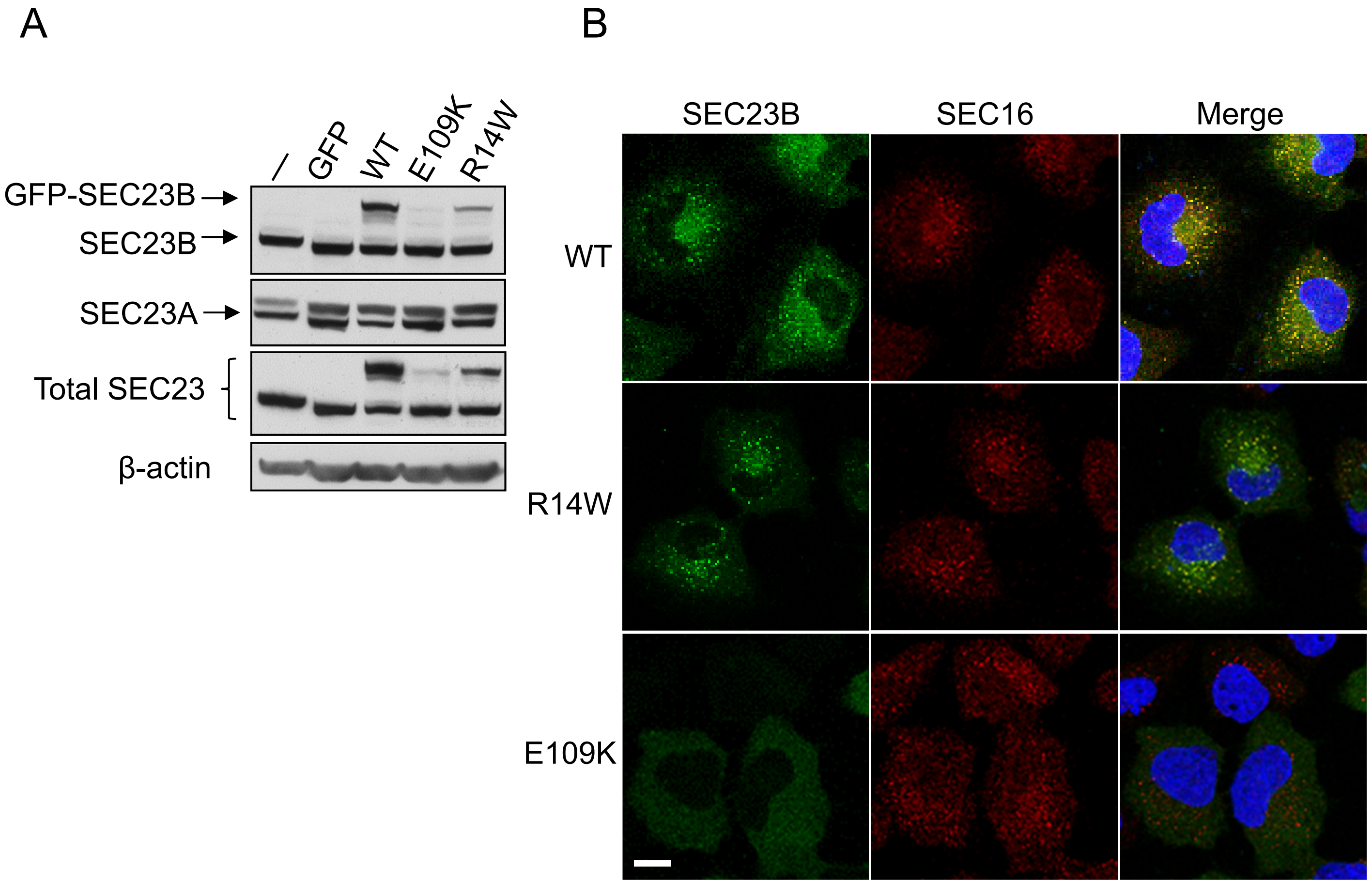


Fig. S5. Decreased protein levels and mis-location of SEC23B result from E109K missense mutation. (A) Western blot analysis was performed to detect protein levels of endogenous and exogenous SEC23B, its endogenous paralog SEC23A and total SEC23 in Nthy-ori 3-1 stable cell lines with GFP tagged WT SEC23B, SEC23B$^{E109K}$ mutant or SEC23B$^{R14W}$ mutant. Experiments were repeated 3 times.

(B) Immunofluorescence staining of Nthy-ori 3-1 stable cell lines was performed to detect the intracellular localization of SEC23B. Nthy-ori 3-1 cells were stained with rabbit anti-Sec16A (red) for ER exit sites and DAPI (blue) for nuclei. Exogenous SEC23B-GFP fusion protein was stained green. Scale bar: 10 μm. Experiments were repeated 3 times.

A

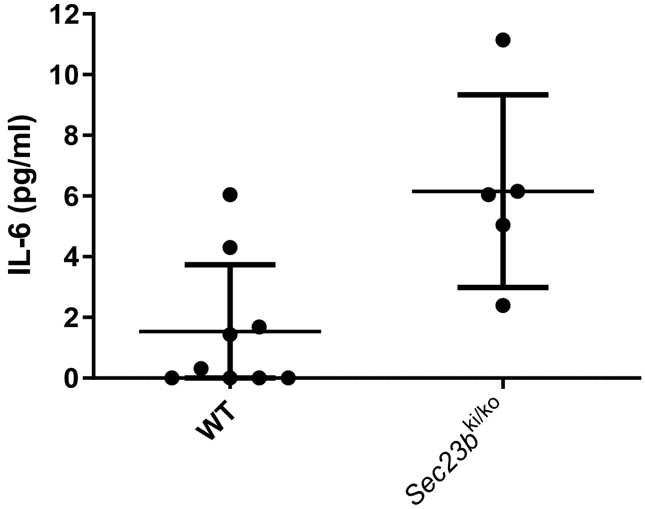


B

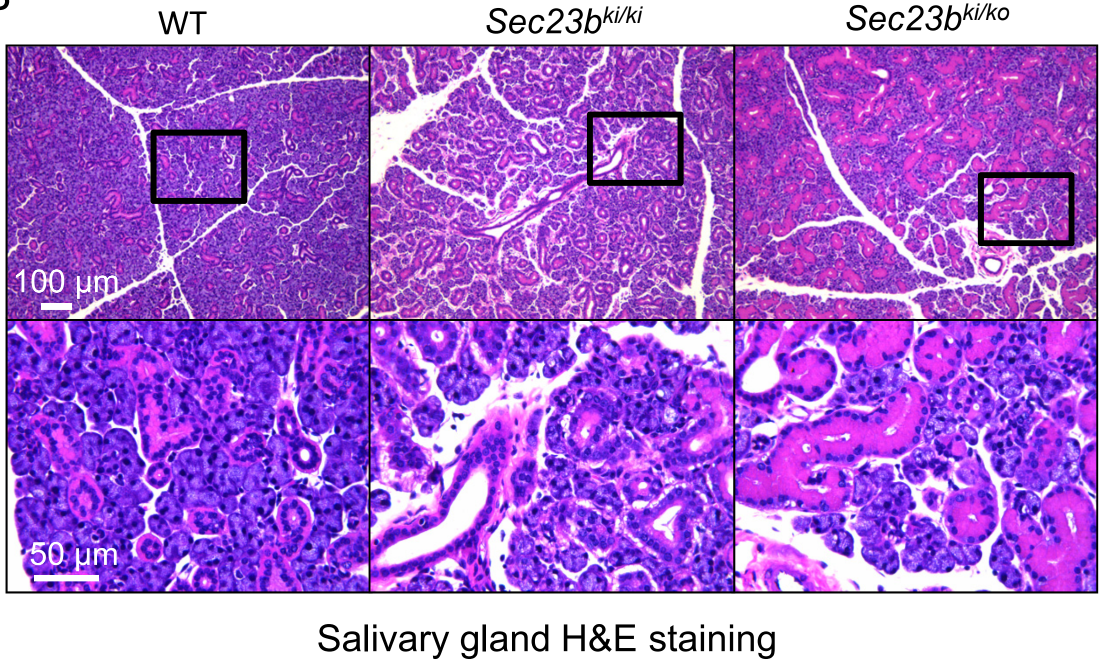


Fig. S6. (A) Comparison of serum IL-6 levels between WT and *Sec23b*$^{ki/ko}$ mice. IL-6 levels were measured using an ELISA kit from Bioassays. (B) No obvious abnormalities in salivary gland (submandibular gland) tissue structures by H&E staining of paraffin-embedded tissues. Scale bars: 100 μm (top) and 50 μm (bottom). Three 2-month old mice from each genotype were analyzed.

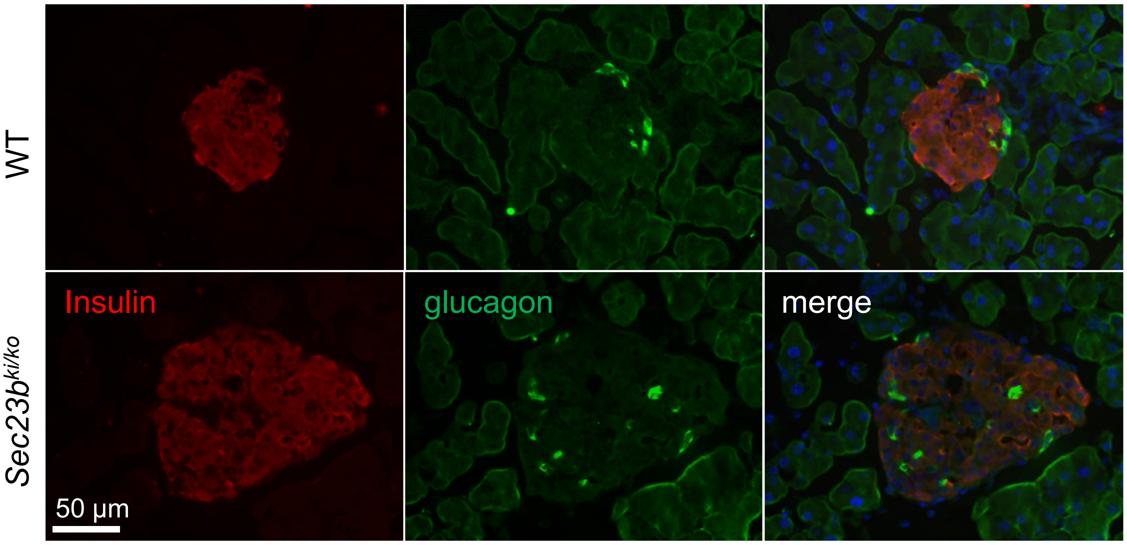


Fig. S7. Normal islet structures in *Sec23b*$^{ko/ki}$ mice. Cryosections of pancreas from WT and *Sec23b*$^{ko/ki}$ mice of 4 month of age were co-stained with mouse anti-glucagon (green) and guinea pig anti-insulin (red) for α- and β-endocrine cells, respectively. Scale bar: 50 μm. Two mice from each genotype were analyzed.
